# Single-cell profiling of CAR-T CD19 cell phenotypes and immune system dynamics in pediatric BCP-ALL

**DOI:** 10.1101/2025.03.27.645594

**Authors:** A. Oszer, B. Pawlik, N. Cwilichowska-Puslecka, P. Marschollek, M. Richert-Przygonska, M. Mielcarek-Siedziuk, JC. Crawford, M. Mazurek, A. Koladiya, K. Kałwak, J. Styczynski, S. Janczar, M. Poreba, MP. Velasquez, KL Davis, W. Mlynarski

## Abstract

**Background:** Chimeric Antigen Receptor T-cell (CAR-T) therapy targeting CD19 has transformed the treatment of relapsed/refractory B-cell precursor acute lymphoblastic leukemia (BCP-ALL). However, the phenotypic heterogeneity of CAR+ T-cells and their interactions with the immune system remain poorly understood. Here, we characterize CAR+ T-cell subsets and persistence in patients receiving standard-of-care tisagenlecleucel in an effort to optimize therapeutic efficacy in pediatric BCP-ALL.

**Methods:** Nineteen pediatric patients with relapsed/refractory BCP-ALL treated with tisagenlecleucel in Poland were included in the study. CAR+ T-cell composition and the peripheral blood mononuclear cell (PBMC) immune system were assessed using mass cytometry, with qPCR validation of CAR+ T-cells. Machine learning algorithms in R classified phenotypes. Statistical analyses examined associations between CAR+ T-cell subsets, immune system changes, and CAR-T therapy outcomes, including cytokine release syndrome (CRS).

**Results:** Infusion product analysis showed a predominance of CD4+ Central Memory (16.6%), MAIT/NKT (7.04%), and Treg memory (63.3%) cells, with high consistency between paired products. Post-infusion, CAR+ Treg memory cells declined to <1%, while CD8+ subsets expanded. PBMC immune system analysis revealed increased monocyte and NK cell counts post-infusion. Patients who experienced CRS had fewer classical monocytes at day 7 and fewer transitional monocytes at day 28. Early and late NK cells increased post-infusion in all patients. However, pre-infusion levels were lower in patients experiencing CRS. CRS correlated with higher CAR+ CD8+ Terminal Effector and MAIT/NKT frequencies in infusion products. While CAR+ composition was not linked to disease burden, prior treatments, or therapy outcomes, changes in monocyte and NK cell dynamics were associated with disease burden, relapse, and CRS.

**Conclusions:** This study provides a detailed characterization of CAR-T CD19 cell composition and post-infusion dynamics in pediatric BCP-ALL patients treated with tisagenlecleucel. Infusion products were predominantly CAR+ CD4+ T cells with a Central Memory phenotype, including a notable Treg memory subset that declined post-infusion. CAR+ CD8+ subsets expanded, coinciding with immune system shifts. These changes correlated with clinical responses, including CRS, highlighting the role of the immune system in CAR-T therapy outcomes and informing strategies to optimize treatment.

**Summary box:** *WHAT IS ALREADY KNOWN ON THIS TOPIC:* CD19-targeted CAR-T therapy has revolutionized the treatment of relapsed/refractory pediatric BCP-ALL. While studies have examined the diverse composition of CAR-T infusion products in adult large B-cell lymphoma (LBCL), there is limited data in the pediatric BCP-ALL setting.

*WHAT THIS STUDY ADDS:* This study demonstrates that standard-of-care CAR-T infusion products (tisagenlecleucel) are primarily composed of CAR+ CD4+ Central Memory and regulatory T cells, which decline post-infusion as CAR+ CD8+ subsets expand. Additionally, monocyte and NK cell dynamics are closely linked to cytokine release syndrome (CRS) and treatment outcomes, emphasizing the critical role of immune system interactions in CAR-T therapy efficacy.

*HOW THIS STUDY MIGHT AFFECT RESEARCH, PRACTICE, OR POLICY:* These findings indicate that monitoring immune system shifts could enhance CRS risk assessment and improve patient stratification, leading to more personalized CAR-T therapy. Additionally, insights into CAR+ T-cell composition may guide the development of strategies to optimize CAR-T efficacy, potentially improving long-term treatment outcomes in pediatric BCP-ALL.

## Background

Chimeric antigen receptor T-cell (CAR-T) therapy targeting CD19 has significantly advanced the treatment of B-cell precursor acute lymphoblastic leukemia (BCP-ALL), the most common malignancy in children(1). It has demonstrated remarkable efficacy in relapsed or refractory B-cell malignancies, including BCP-ALL(2) and diffuse large B cell lymphoma (DLBCL)(3). Tisagenlecleucel (tisa-cel, Kymriah; Novartis Pharmaceutical), a CD19-directed, genetically modified, autologous T-cell therapy, is currently the only CAR-T CD19 therapy approved by both the U.S. Food and Drug Administration (FDA)(4) and the European Medicines Agency (EMA)(5) for pediatric BCP-ALL. This therapy has significantly reduced mortality after relapse in high-risk patients. Clinical data indicate that three-year event-free survival (EFS) rates post-CAR T therapy is 44%, while the overall survival (OS) rate is 63%(6), an improvement when compared to historical controls prior to CAR-T CD19 therapy, where five-year OS was 52%(7).

Significant heterogeneity exists in CAR+ T-cell composition within infusion products (IP), particularly in LBCL(8, 9). However, it remains unclear whether these phenotypic variations significantly impact clinical outcomes, as ongoing research continues to explore this aspect(9–11). Further, in-depth characterization of tisagenlecleucel products and their post-infusion behavior in children with BCP-ALL has not been previously studied. Emerging evidence suggests that the persistence of CAR+ T-cells in peripheral blood post-infusion may play a crucial role in determining therapy outcomes (12–14).

How pre-infusion tisa-cel composition influences post-infusion CAR+ T-cell subsets, disease response and toxicity in the pediatric population remains unknown. To address this gap, we analyzed a cohort of 19 BCP-ALL patients receiving standard-of-care CAR-T therapy with tisagenlecleucel. Using mass cytometry, we characterized the phenotype of CAR+ T-cells in infusion products and tracked their persistence in peripheral blood on days 7 and 28. Our results highlight that the CAR+ Treg subset was dominant in the infusion product but was absent on days 7 and 28, where CAR+ CD8+ subsets increased and became dominant. Additionally, we profiled T cells in peripheral blood at the time of apheresis and on day 0 before CAR-T CD19 infusion. We also assessed changes in non-CAR T mononuclear cells, particularly monocytes and NK cells, which increased during the first month following CAR-T infusion. Notably, their association with cytokine release syndrome (CRS), a common side effect affecting over 70% of recipients(6) was of particular interest. Further, we examined relationships between clinical outcomes, pre-treatment patient status, and prior therapies—factors that have not been comprehensively studied before and appear to vary independently of CAR+ T-cell subsets in the infusion product. Our findings offer valuable insights to optimize therapeutic efficacy and mitigate treatment-associated toxicities, ultimately improving clinical outcomes for pediatric patients.

## Methods

### Study cohort

Patients aged 2-16 years old with relapsed/refractory BCP-ALL who received tisagenlecleucel as a standard-of-care treatment in Poland. All patients provided informed consent, and the study was approved by the Human Research Ethics Committee of the Medical University of Lodz (approval no. RNN/93/22/KE). Clinical data were obtained retrospectively from the medical record. Patients were observed between September 15, 2022, and August 21, 2024, with follow-up until February 12, 2025. Treatment response was assessed using PCR-based minimal residual disease (PCR-MRD) in bone marrow biopsies, and toxicity was evaluated using the Penn scale for CRS(15) and the ICANS scale for neurotoxicity(16).

This study was conducted in accordance with the STROBE (Strengthening the Reporting of Observational Studies in Epidemiology) guidelines for cohort studies to ensure transparent and comprehensive reporting(17).

### Mass cytometry

The Maxpar Direct Immunoprofiling Assay (MDIPA) antibody panel (Standard BioTools, USA) was extended with four additional antibodies: anti-human CD279/PD-1 (EH12.2H7)/165Ho (Fluidigm, USA), CTLA-4 (Bio-Techne, USA), CD274/PD-L1 (29E.2A3)/175Lu (Fluidigm, USA), and anti-FMC63 scFv (Y45). Anti-FMC63 and CTLA-4 were labeled in-house with 142Nd and 169Tm using the Maxpar X8 kit (Supplementary Table 1). CAR+ expressing were defined by anti-FMC63 expression on CD3+ cells.

Infusion products (tisagenlecleucel) were collected post-administration, stored at 4°C and processed within 24 hours. PBMCs were collected at apheresis, day 0 (pre-infusion), and days 7 and 28 post-infusion. Samples were stored at 4°C and processed within 24 hours.

Samples processing followed a modified protocol (18). CAR-T cells were diluted in 50 mL PBS (without ions), vortexed, centrifuged (400 × g, 5 min) and pellet was resuspended in 1 mL Cell Staining Buffer (CSB). Whole blood was treated with RBC buffer, washed with PBS, centrifuged, and resuspended in CSB. Approximately 2 × 10⁶ cells were incubated with antibodies cocktail for 30 min at room temperature (RT) and diluted to 5 mL with CSB. After centrifugation at 400 × g for 5 min, the pellet was resuspended in 1 mL 1.6% formaldehyde and incubated for 10 min at RT. The sample was centrifuged at 800–900 × g for 5 min, and the supernatant was carefully removed. The pellet was resuspended in 1 mL Fix and Perm Buffer (Standard BioTools, USA) containing 125 nM Cell-ID Intercalator-Ir (Standard BioTools, USA) and incubated overnight at 4°C. The sample was centrifuged at 800–900 × g for 5 min, 90% of the supernatant was removed, and the pellet was stored at −80°C until acquisition.

Samples were acquired using a Helios® mass cytometer (Standard BioTools, USA) at a rate of 250– 350 events per second. Data were stored in .fcs format, and interrupted measurements were concatenated using CyTOF Software v7.0 (Standard BioTools). PBMCs from healthy children (n=4) contained approximately 300,000 cells per sample, while patient samples (n=77) ranged from 100,000 to 500,000 cells per sample.

### qPCR analysis of CAR-T cells

Genomic DNA was extracted from patient samples using the Invisorb Spin Universal Kit (Invitek Diagnostics, Germany) and quantified by UV spectroscopy (NanoDrop 8000, Life Technologies, USA). DNA samples were diluted to 20 ng/µL in nuclease-free water. PBMC samples were collected on day 7 (n=17) and day 28 (n=16) post-CAR-T CD19 infusion. Transgene copies were quantified by qPCR using custom-designed primers/probes and the TaqMan Gene Expression Master Mix (Applied Biosystems, USA). The assay targeted the CAR T transgene with the RNaseP Copy Number Reference Assay (Applied Biosystems, USA) as an internal control. qPCR was performed on the AriaMx Real-Time PCR System (Agilent, USA) with an initial activation (50°C, 2 min), denaturation (95°C, 10 min), and 45 cycles of 95°C (15 sec) and 60°C (1 min). Negative controls (healthy donor DNA and NTCs) ensured specificity. CAR T transgene levels were calculated using the 2−ΔCt method(19).

### Statistical analysis

Raw FCS files were bead-normalized(20) and transformed using the inverse hyperbolic sine (arsinh) function (cofactor = 5). Data were cleaned in OMIQ by gating out beads and filtering based on residual, center, offset, width, event length, and DNA intercalator signals in relation to time using Gaussian discrimination(21).

PBMCs from healthy children (n=4, ∼300,000 cells/sample) were manually gated using the gating strategy in Supplementary Figure 1A, 1B. A classification model was built in R, based on Mahalanobis distance, to identify all cell populations with a focus on T cells (Supplementary Figure 2A). Cross-validation was performed using healthy control samples (Supplementary Figure 2B, C).

Patient samples (n=77, ∼100,000–500,000 cells per sample) were gated for T cells (CD45⁺CD3⁺), with CAR⁺ and CAR⁻ T cells distinguished by CAR expression. A supervised classification method was applied in R, first classifying all cell populations, then non-CAR T cells, and finally CAR+ T cells (Supplementary Figure 2D). Median protein expression of CAR+T-cells was analyzed in OMIQ after subsampling to 1,000 CAR+ T-cells per patient. Classified FCS files were manually checked for single-cell data quality (Supplementary Figure 3).

Percentages and counts of classified populations were exported to GraphPad Prism (v10, GraphPad Software, USA). Figures were generated using GraphPad Prism and BioRender (https://biorender.com). Data analysis was performed in R (http://www.r-project.org) and GraphPad Prism. t-SNE was used to visualize time-course data and infusion products (subsampled to 10,000 CAR+ T-cells per patient) based on classification markers including CD45, CCR6, IL-3R, CD4, CD8a, CD11c, CD45RO, CD45RA, CD161, CCR4, IL-2Ra, CD27, CXCR3, CXCR5, CD28, TCR γδ, CCR7, CD14, CD3, CD66b, IL-7Ra and classified cells. Additionally, t-SNE was applied to visualize CAR+ T-cells in IP, on day 7 and 28 post-infusion, using the same markers, to assess expression of activation (IL-2Ra(CD25), CD28, CD38, HLA-DR, IL-7Ra(CD127)) and exhaustion (CTLA-4, PD-1(CD279), PD-L1(CD274), CRTH2(CD294)) T-cells markers.

CAR+ T-cell subsets and other PBMC components were compared using either a paired t-test or a Wilcoxon signed-rank test adjusted for multiple comparisons using the FDR method, depending on the results of the Shapiro-Wilk test for normality of the data. The Mann-Whitney U test assessed differences between groups, and linear regression validated CAR⁺ gating by qPCR. Overall survival (OS) and event-free survival (EFS) were estimated using the Kaplan–Meier method.

## Results

### Patient Characteristics

We analyzed 19 consecutive patients with relapsed/refractory (R/R) BCP-ALL who received commercial tisagenlecleucel (tisa-cel) therapy in Poland between September 15, 2022, and August 21, 2024, with follow-up until February 12, 2025 (Table 1). At the time of tisa-cel infusion, eight patients (42%) had positive measurable disease, as detected by PCR-based minimal residual disease (PCR-MRD) assessment in the bone marrow (<1×10⁻⁴: negative; ≥1×10⁻⁴: positive). Among these, only one patient had a high disease burden (88% bone marrow blasts) prior to infusion, whereas the remaining PCR-MRD-positive patients had less than 1% blasts by morphology. Fourteen patients (74%) received CAR-T therapy for first relapse, either relapse after allogeneic hematopoietic stem cell transplantation (allo-HSCT) or after failure to achieve complete remission (CR) after one cycle of re-induction therapy for relapse. Four patients were treated for second relapse and one patient for fourth relapse. Prior to CAR-T therapy, nine patients (47%) had received allo-HSCT, two had received blinatumomab, and six had received with inotuzumab ozogamicin. Cytokine release syndrome occurred in 74% of patients, with a maximum severity of grade 2. Seven patients received tocilizumab, all within the first seven days post-infusion. Immune effector cell-associated neurotoxicity syndrome (ICANS) developed later than CRS, with CRS severity peaking between days 0–6 post-infusion and ICANS severity peaking between days 3–8. ICANS was observed in three patients, with two experiencing grade 2 neurotoxicity and one experiencing grade 4 neurotoxicity. Complete remission with sustained B-cell aplasia was achieved in 14 patients (73%); however, four subsequently relapsed within one year. One patient experienced disease progression, and two patients died due to leukemia relapse or resistance during follow-up, with a median follow-up of one year (Supplementary Figure 4).

**Table 1.**
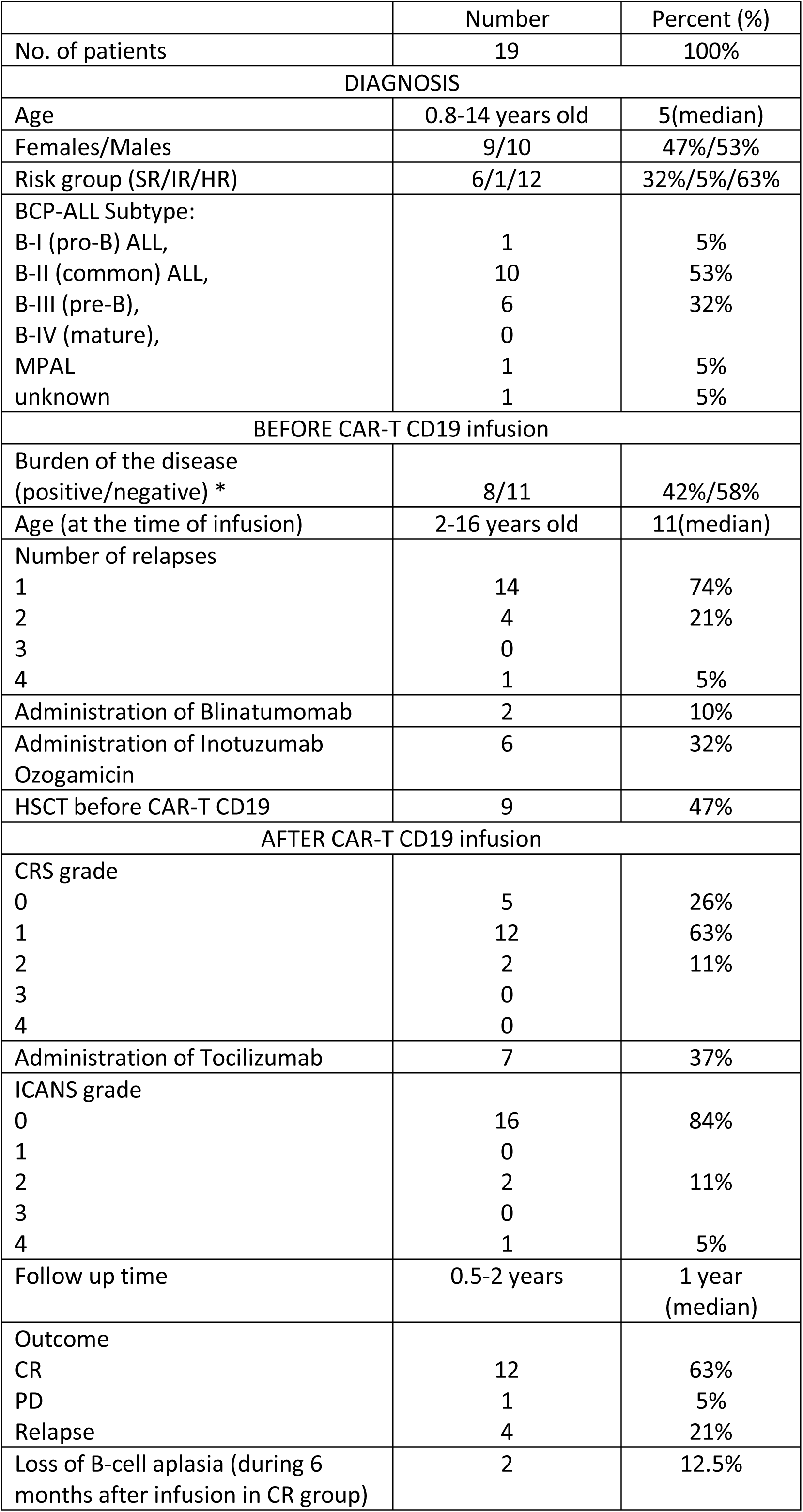
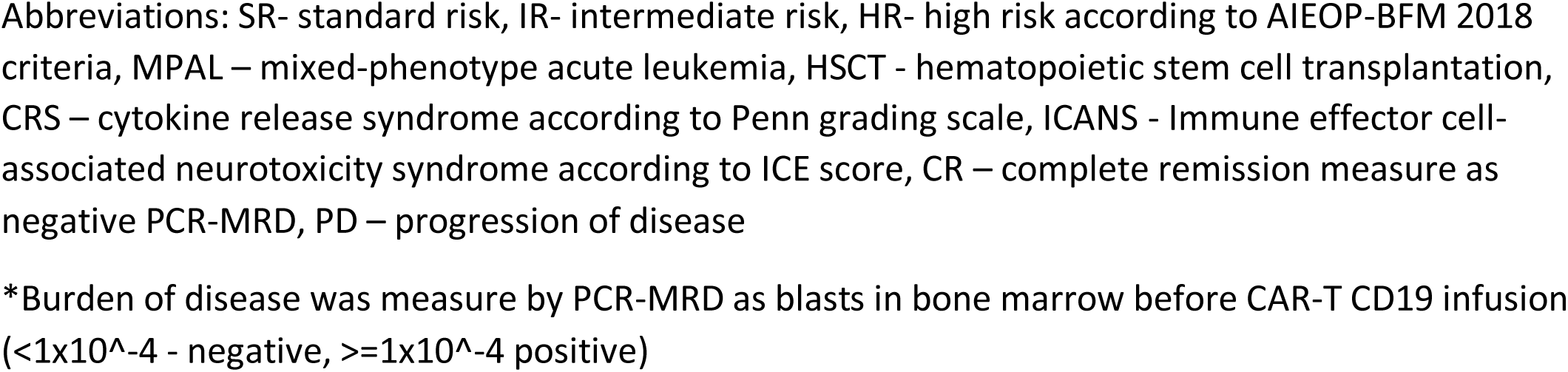
Patient characteristics.

|  | Number | Percent (%) |
| --- | --- | --- |
| No. of patients | 19 | 100% |
| **DIAGNOSIS** |  |  |
| Age | 0.8-14 years old | 5(median) |
| Females/Males | 9/10 | 47%/53% |
| Risk group (SR/IR/HR) | 6/1/12 | 32%/5%/63% |
| BCP-ALL Subtype: |  |  |
| B-I (pro-B) ALL, | 1 | 5% |
| B-II (common) ALL, | 10 | 53% |
| B-III (pre-B), | 6 | 32% |
| B-IV (mature), | 0 |  |
| MPAL | 1 | 5% |
| unknown | 1 | 5% |
| **BEFORE CAR-T CD19 infusion** |  |  |
| Burden of the disease<br>(positive/negative) * | 8/11 | 42%/58% |
| Age (at the time of infusion) | 2-16 years old | 11(median) |
| Number of relapses |  |  |
| 1 | 14 | 74% |
| 2 | 4 | 21% |
| 3 | 0 |  |
| 4 | 1 | 5% |
| Administration of Blinatumomab | 2 | 10% |
| Administration of Inotuzumab<br>Ozogamicin | 6 | 32% |
| HSCT before CAR-T CD19 | 9 | 47% |
| **AFTER CAR-T CD19 infusion** |  |  |
| CRS grade |  |  |
| 0 | 5 | 26% |
| 1 | 12 | 63% |
| 2 | 2 | 11% |
| 3 | 0 |  |
| 4 | 0 |  |
| Administration of Tocilizumab | 7 | 37% |
| ICANS grade |  |  |
| 0 | 16 | 84% |
| 1 | 0 |  |
| 2 | 2 | 11% |
| 3 | 0 |  |
| 4 | 1 | 5% |
| Follow up time | 0.5-2 years | 1 year<br>(median) |
| Outcome |  |  |
| CR | 12 | 63% |
| PD | 1 | 5% |
| Relapse | 4 | 21% |
| Loss of B-cell aplasia (during 6<br>months after infusion in CR group) | 2 | 12.5% |

Abbreviations: SR- standard risk, IR- intermediate risk, HR- high risk according to AIEOP-BFM 2018 criteria, MPAL – mixed-phenotype acute leukemia, HSCT - hematopoietic stem cell transplantation, CRS – cytokine release syndrome according to Penn grading scale, ICANS - Immune effector cell-associated neurotoxicity syndrome according to ICE score, CR – complete remission measure as negative PCR-MRD, PD – progression of disease
\*Burden of disease was measure by PCR-MRD as blasts in bone marrow before CAR-T CD19 infusion ( $<1 \times 10^{-4}$ - negative, $\geq 1 \times 10^{-4}$ positive)

### Phenotypic profiling of CAR+ T-cells in tisa-cel infusion products

Nineteen patients contributed 77 samples across several timepoints as shown in Figure 1A. Composition of cell subtypes in peripheral blood at apheresis (n=5), day 0 (n=18), day 7 (n=19) and day 28 (n=16) (Figure 1B) and Tisa-cel infusion products (n=19) (Figure 1C). Tisa-cell products demonstrated high viability of CD3+ T cells (98.7% median; range 94.3%–99.6%) (Figure 1D), with CAR+ T-cells constituting 17.6% (median; range 7.91%–42.9%) of CD3+ cells (Figure 1E, Supplementary Figure 5A). CAR+ T-cell subsets were further characterized. Among CD8+ CAR+ T-cells, Central Memory (CCR7hi, CD45RA-, CD45RO+) cells represented 1.62% (median; range 0.01%–9.31%), Effector Memory (CCR7lo/-, CD27+) 0.08% (median; range 0%–1.78%), and Terminal Effector (CCR7lo/-, CD27-) 0.35% (median; range 0.01%– 4.08%) in relation to the total CAR+ T-cell population (Figure 1F). Among CD4+ CAR+ T-cells, Central Memory comprised 16.6% (median; range 1.35%–55.4%), Effector Memory 1.97% (median; range 0.11%–9.02%), and Terminal Effector 0.5% (median; range 0.04%–13.7%) of total CAR+ T-cells (Figure 1G). Notably, Treg memory (CXCR5-, CD45RA-, CD25hi, CD127lo/-) predominated, accounting for 63.3% of CAR+ T-cells in tisa-cel products (median; range 30.3%–88.7%) (Figure 1H, Supplementary Figure 5B, C). Th1-, Th2-, and Th17-like cells were rare (<0.5%) (Figure 1I). CAR+ γδ T cells and MAIT/NKT cells comprised 0.27% and 7.04%, respectively (Figure 1H). No CAR+ Naïve CD4+ or CD8+ cells were detected. Overall, tisa-cel infusion products were primarily composed of MAIT/NKT (7.04%), CD4+ Central Memory (16.6%), and predominantly Treg memory (63.3%) cells, while other subsets comprised less than 2% of CAR+ T-cells. Three patients received two infusion products (IP A and IP B) derived from a single apheresis. Linear regression analysis of CAR+ T-cell subsets between paired products demonstrated a strong correlation (R² = 0.9894, p < 0.0001) (Figure 1J, Supplementary Figure 6), indicating high product consistency within patients.

**Figure 1.**
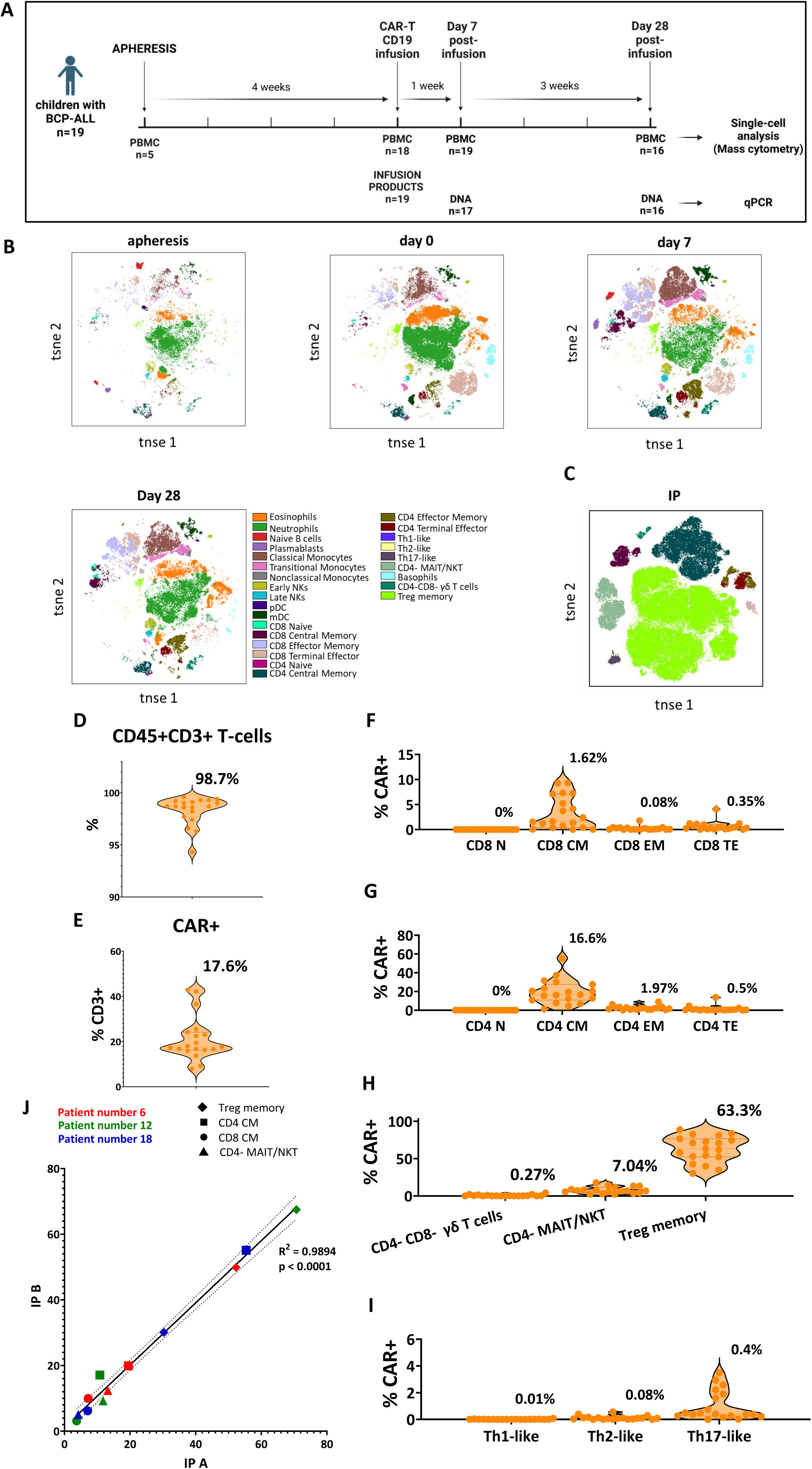
Phenotypic diversity in CAR+ T-cells within CAR-T CD19 infusion products (tisa-cel). **A.** Schematic of sample collection and analysis, including mass cytometry and qPCR from pediatric BCP-ALL patients (Supplementary Tables 1, Supplementary Figure 1). Figure created using BioRender.com. **B.** Phenotypic distribution of cell subsets based on timepoint classified using Mahalanobis distance (Supplementary Figures 2–3). **C.** T cell subsets within CAR+ T-cells in infusion products. **D.** Proportion of CD3+CD45+ T-cells in infusion products (median 98.7%, range 94.3-99.6%). **E.** Proportion of CAR+ cells within CD3+ cells (median 17.6%, range 7.91-42.9%). **F.** Subpopulations of CD8+ CAR+ in Naïve, Central Memory, Effector Memory, and Terminal Effector subsets. **G.** Subpopulations of CD4+ CAR+ in Naïve, Central Memory, Effector Memory, and Terminal Effector subsets. **H.** γδ T cells (median 0.27%, range 0.03-4.17%), CD4-MAIT/NKT cells (median 7.04%, range 2.33-17.8%), and CAR+ Tregs (median 63.3%, range 30.3-88.7%) as key components of the infusion product. **I.** CAR+ Th cell subsets. **J.** High correlation of per-patient infusion products when released as two individual products (R² = 0.9894, p < 0.0001, Supplementary Figure 6).

### Dynamics of CAR+ T-cell subpopulations

Mass cytometry and qPCR quantification showed high concordance at day 7 (R² = 0.9043, p < 0.0001) and day 28 (R² = 0.9825, p < 0.0001) post-infusion (Figure 2A). Comparing PBMC at time of T cell apheresis (n=5) to manufactured tisa-cel products revealed a higher proportion of CD8 Naïve, CD8 Effector Memory, and CD4 Effector Memory T cells (p<0.05) in the apheresis PBMC and a significantly lower fraction of Treg memory cells (p<0.01) (Figure 2B). No other significant differences were observed (Supplementary Figure 7).

**Figure 2.**
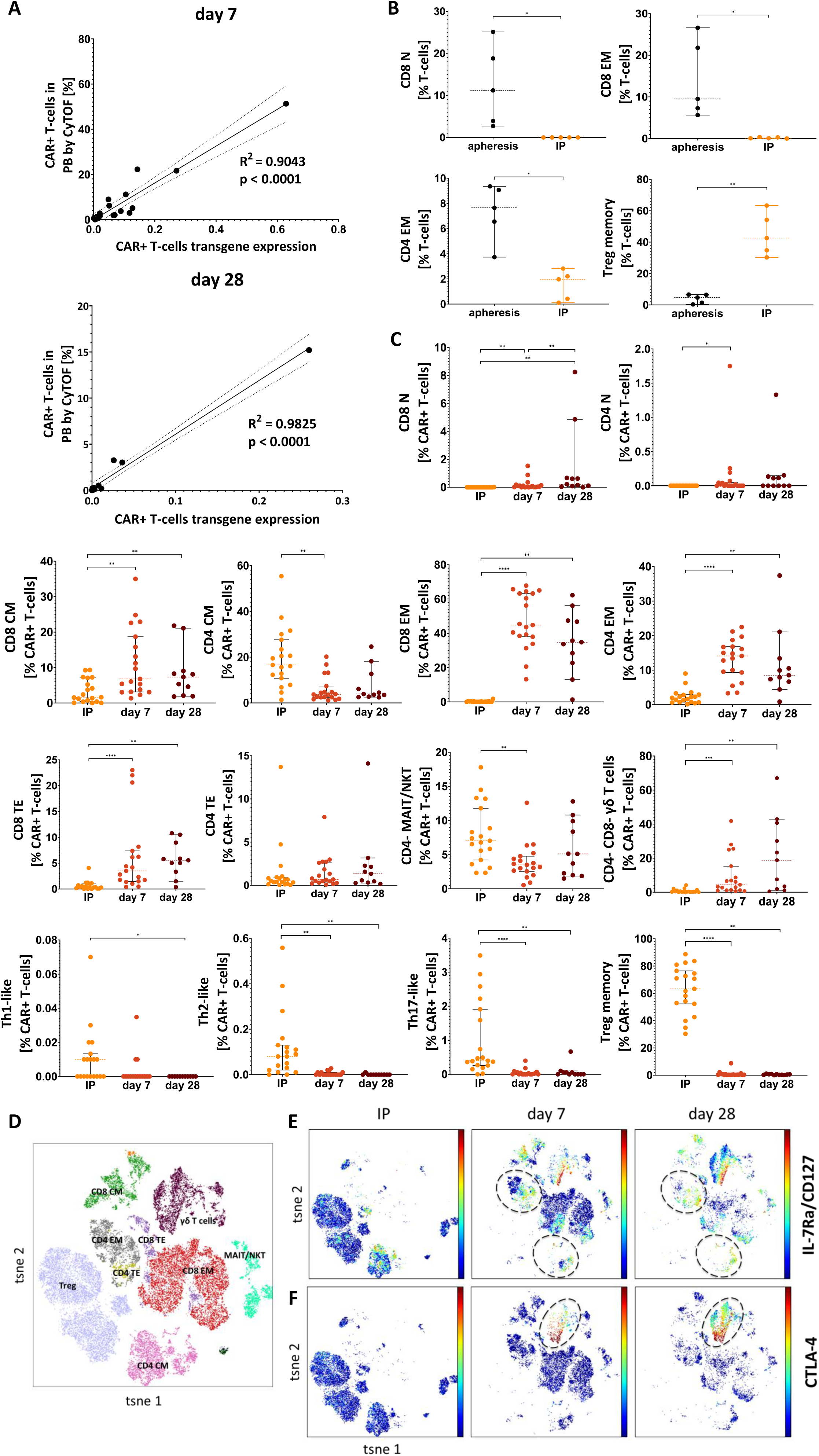
In vivo dynamics of CAR+ T-cell subpopulations after infusion. **A.** Correlation between CAR+ cell percentages in PBMCs by mass cytometry and CAR+ transgene expression at day 7 (R² = 0.9043, p < 0.0001) and day 28 (R² = 0.9825, p < 0.0001). **B.** Comparison of T-cell subpopulations in PBMCs at apheresis vs. CAR+ T-cells in infusion products, (*p < 0.05; *\*p < 0.01, paired t-test, Supplementary Figure 7*). **C.** Longitudinal analysis of CAR+ T cell subsets from infusion to day 7 and day 28 (*p < 0.05; **p < 0.01; ***p < 0.001; *\*\*\*p < 0.0001, Wilcoxon test, FDR-adjusted*). **D.** CAR+ T-cell subsets at all timepoints classified using Mahalanobis distance (Supplementary Figure 8a). **E.** IL-7Rα (CD127) expression on CAR+ T-cells over time (infusion, day 7, and day 28), with additional activation markers in Supplementary Figures 8b and 8c. **F.** CTLA-4 expression on CAR+ T-cells over time (infusion, day 7, and day 28), with additional data in Supplementary Figure 8d.

CAR+ Treg memory cells, which were prominent in the tisa-cel products, decreased significantly by day 7 (<0.5%, p<0.0001) and day 28 (p<0.001) compared to infusion products (IP). Other CAR+ T-cell subsets also declined, including CD4 Central Memory (day 7: p<0.001; day 28: p<0.05), CD4- MAIT/NKT (day 7: p<0.01), Th1-like (day 7: p<0.05), Th2-like (day 7: p<0.0001; day 28: p<0.01), and Th17-like cells (day 7: p<0.0001; day 28: p<0.01) (Figure 2C).

Conversely, the proportion of CD8 Naïve cells increased post-infusion (day 7: p<0.0001; day 28: p<0.01), with further expansion between day 7 and 28 (p<0.01). Similar trends were observed for CD8 Central Memory (day 7: p<0.01; day 28: p<0.05), CD8 Effector Memory (day 7: p<0.0001; day 28: p<0.001), and CD8 Terminal Effector cells (day 7: p<0.0001; day 28: p<0.001). CD4 Naïve, CD4 Effector Memory, and CD4-CD8- γδ T cells also increased (p<0.01), with further expansion between day 7 and 28 (p<0.05). No significant differences were observed for CD4 Terminal Effector cells (Figure 2C).

t-SNE visualization of CAR+ T-cells (Figure 2D) demonstrated distinct clustering patterns over time. Further analysis of activation and exhaustion markers in CAR+ T cells revealed an increased expression of IL-7Rα (CD127) on days 7 and 28 post-infusion compared to the infusion product (IP), particularly in CAR+ CD4+ Central Memory and Effector Memory subsets (Figure 2E). Additionally, CTLA-4 expression was elevated on days 7 and 28 post-infusion compared to IP, especially in CAR+ CD4−CD8− γδ T cells (Figure 2F). Changes in other activation and exhaustion markers in CAR+ T cells are presented in Supplementary Figure 8A-D.

### Immune system dynamics

Given that CD19-targeted CAR-T therapy has been shown to influence the immune system in other diseases(22), we investigated changes in the composition of CAR- leukocytes during treatment in this cohort. At apheresis, only Late NK cells have are significantly different between PBMCs collected at apheresis and those at day 0 pre-infusion (p<0.05) (Supplementary Figure 9A-B).

To further characterize immune system changes, we analyzed PBMCs from 16 patients at day 0 (pre-infusion), day 7, and day 28 post-infusion. As expected, we observed a significant decrease in Naïve B cells, consistent with expected B-cell aplasia (day 0–7: p<0.01; day 0–28: p<0.05). Neutrophil counts decreased significantly within 7 days post-infusion (p<0.001) but rebounded by day 28 (p<0.05).

Plasmacytoid dendritic cells (pDCs) increased within 7 days (p<0.05), whereas myeloid dendritic cells (mDCs) declined between day 7 and 28 (p<0.05). Monocytes, as well as Early and Late NK cells, significantly increased within 7 days post-infusion (p<0.01, p<0.05, p<0.05, respectively) and remained elevated at day 28 compared to day 0 (p<0.01, p<0.001, p<0.01, respectively) (Figure 3A-D, Supplementary Figure 9C).

**Figure 3.**
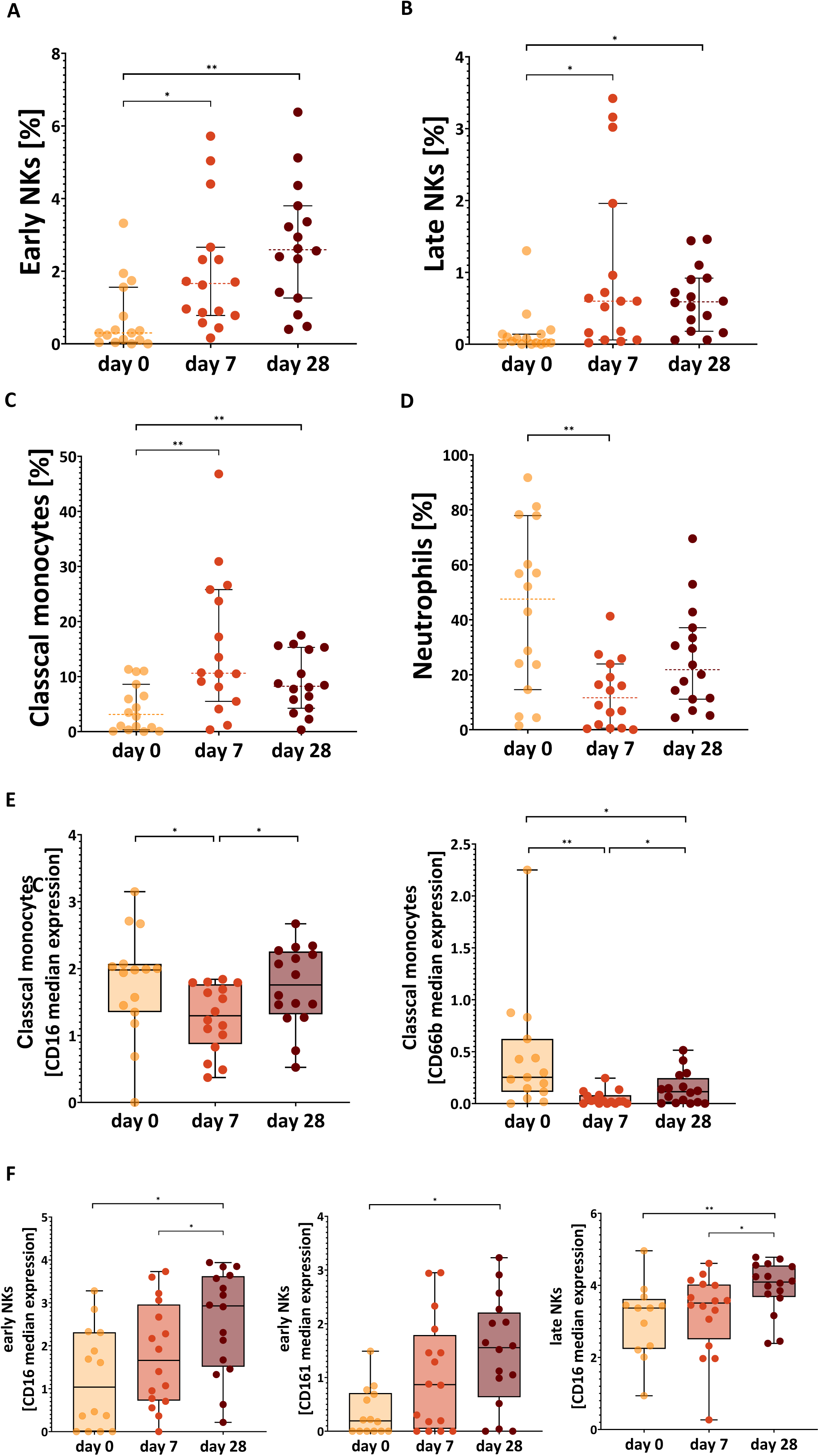
Immune system dynamics. **A.** Longitudinal analysis (day 0, 7, 28) of early/late NK cells, classical monocytes, pDCs, and mDCs, along with B-cell aplasia and a decrease in neutrophils at day 7 (Supplementary Figure 9b, *p < 0.05; **p < 0.01; ***p < 0.001). **B.** CD16 and CD66b expression on classical monocytes decreases at day 7, followed by recovery at day 28 (*p < 0.05; **p < 0.01; ***p < 0.001). **C.** CD16 and CD161 expression on early NK cells increases at day 28, while late NK cells show higher CD16 at day 28 compared to day 0 (*p < 0.05; **p < 0.01; ***p < 0.001).

Despite the observed increase in monocyte counts, the expression of activation markers (CD16, CD66b) on monocytes decreased significantly by day 7 (p<0.01, p<0.001) before partially recovering by day 28 (p<0.05, p<0.05) (Figure 3E). In contrast, activation markers on Early and Late NK cells increased over time. CD16 expression on NK cells rose between day 0 and 28 (p<0.05) and between day 7 and 28 (p<0.05). Additionally, CD161 expression on Early NK cells increased between day 0 and 28 (p<0.05) (Figure 3F).

### Association with burden of disease, relapse and CRS

Despite interpatient variability in the composition of CAR+ T-cell subsets within infusion products, no associations were observed between CAR+ T-cell subset composition and patient risk group, disease burden, prior treatment with blinatumomab, inotuzumab ozogamicin, or HSCT. Furthermore, CAR+ T-cell composition was not correlated with treatment outcomes or the development of ICANS (Table 2). However, a higher percentage of CAR+ CD4+ Effector Memory was observed in females compared to males within IP, as well as in PBMCs at day 28 post-infusion (Supplementary Figure 10A-B). Additionally, age-associated differences were noted in CAR+ CD8+ Effector Memory T, eosinophils, and late NK cells in PBMCs post-infusion (Supplementary Figure 11A-C).

**Table 2.** Influence of various factors during therapy on CAR+ T-cell subpopulations within the infusion product (tisa-cel). P-values were determined using the Mann-Whitney U test. Significant p value is highlighted in red in the table. Refer to Supplementary Figure 10 for violin plots illustrating significant differences in CAR+ CD4 Effector Memory between males and females within the infusion product and in PBMCs on day 7 post-infusion. Refer to Supplementary Figure 11 for violin plots illustrating significant differences in CAR+ CD8 Effector Memory between patients younger than 10 years and older than 10 years in PBMC on day 7 post-infusion.

| Variable | CAR+ T-cells in CAR-T CD19 infusion product |  |  |  |  |  |  |  |  |
| --- | --- | --- | --- | --- | --- | --- | --- | --- | --- |
| | CD8 Central Memory | CD8 Effector Memory | CD8 Terminal Effector | CD4 Central Memory | CD4 Effector Memory | CD4 Terminal Effector | CD4-MAIT/NKT | CD4-CD8-$\gamma\delta$ T cells | Treg memory |
| Sex male/female 10/9 | p=0.8421 | p=0.9673 | p=0.7050 | p=0.7802 | **p=0.0172** | p=0.1377 | p=0.7802 | p=0.3555 | p=0.9682 |
| Risk group* HR/non-HR 12/7 | p=0.8369 | p=0.3500 | p=0.7892 | p=0.7108 | p>0,9999 | p=0.6340 | p=0.9671 | p=0.1414 | p>0,9999 |
| Blinatumomab** yes/no 2/17 | p>0.9999 | p=0.5673 | p=0.9240 | p=0.8421 | p>0,9999 | p=0.7427 | p=0.8421 | p=0.2924 | p=0.9474 |
| Inotuzumab** yes/no 7/12 | p=0.8314 | p=0.1997 | p=0.7176 | p=0.7012 | p=0.7012 | p=0.5077 | p=0.8983 | p=0.4662 | p=0.7654 |
| HSCT** yes/no 9/10 | p=0.7197 | p=0.7332 | p=0.7950 | p=0.1823 | p=0.3154 | p=0.7377 | p=0.7197 | p=0.4462 | p=0.3154 |
| Burden of the disease*** positive/negative 8/11 | p=0.3511 | p=0.1221 | p= 0.8880 | p=0.2723 | p=0.8404 | p=0.9486 | p=0.6574 | p=0.2051 | p=0.5448 |
| Age at the time of infusion $\geq 10$/$<10$ 13/6 | p=0.6388 | p=0.1997 | p=0.1863 | p=0.6388 | p=0.4670 | p=0.0844 | p=0.6388 | p=0.6384 | p=0.7654 |
| CRS grade $\geq 1$/0 14/5 | p=0.1560 | p=0.0913 | **p=0.0460** | p=0.4998 | p=0.4998 | p=0.9802 | **p=0.0143** | p=0.6213 | p=0.2976 |
| ICANS $\geq 1$/0 3/16 | p=0.9577 | p=0.4076 | p=0.8947 | p=0.7926 | p=0.3034 | p=0.3509 | p=0.1383 | p=0.7905 | p=0.9577 |
| Outcome**** CR/non-CR 14/5 | p=0.2976 | p=0.4851 | p=0.4847 | P=0.8932 | p=0.1068 | p=0.9486 | p=0.4998 | p=0.6213 | p=0.6868 |

\* at the time of diagnosis
\*\* administration before CAR-T CD19 therapy
\*\*\* burden of disease (% of blasts in bone marrow before CAR-T CD19 infusion) – MRD ( $<1 \times 10^{-4}$ - negative, $\geq 1 \times 10^{-4}$ positive)
\*\*\*\* outcome at 6-months after CAR-T CD19 therapy

In contrast, the composition of non-T-cell PBMC subsets was associated with disease burden, relapse, and CRS. Patients with positive PCR-MRD (≥1 × 10⁻⁴ blasts in bone marrow) exhibited fewer Classical Monocytes and Early and Late NK cells at apheresis compared to PCR-MRD-negative patients (p<0.05) (Figure 4A. Additionally, median NCAM expression on Early NK cells was lower in patients with positive PCR-MRD (p<0.01) (Figure 4B). By day 28 post-infusion, PCR-MRD-positive patients had fewer non-classical monocytes (p<0.05) but higher CD66b expression on monocytes compared to PCR-MRD-negative patients (Figure 4C).

**Figure 4.**
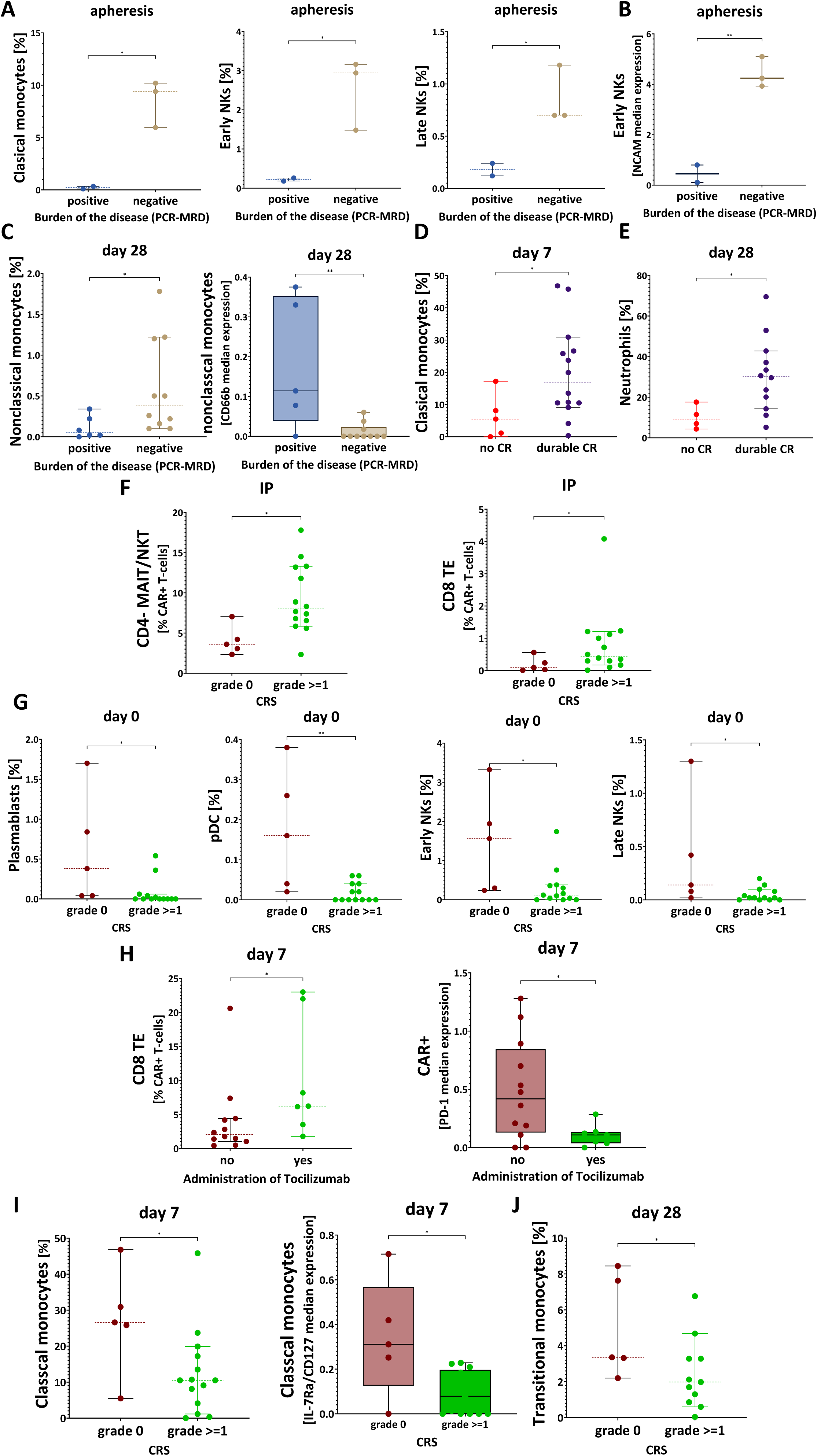
Post-Infusion PBMC Composition and Association with Clinical Characteristics. **A.** Abundance of immune cell subsets at apheresis based on disease burden at infusion (*p < 0.05, Mann-Whitney U test). **B.** Early NK cells in PCR-MRD negative patients show significantly increased NCAM (CD56) expression (**p < 0.01). **C.** Abundance of non-classical monocytes and their expression of CD66b at day 28 based on disease burden at infusion (*p < 0.05; **p < 0.01). **D.** Abundance of classical monocytes at day 7 post-infusion based on response at day 28 to tisa-cel (*p < 0.05), **E.** Abundance of neutrophils at day 28 based on response to tisa-cel at day 28 (*p < 0.05). **F.** Abundance CAR+ CD4⁻ MAIT/NKT cells and CAR+ CD8 Terminal Effector cells based on development of CRS (*p < 0.05). **G.** Pre-infusion PBMC composition at day 0 indicates higher levels of plasmablasts, plasmacytoid dendritic cells (pDCs), early NK cells, and late NK cells in patients who did not develop CRS (*p < 0.05; **p < 0.01). **H.** At day 7 post-infusion, tocilizumab-treated patients exhibit increased CAR+ CD8 Terminal Effector cells and significantly lower PD-1 expression across all CAR+ T cells (*p < 0.05). **I.** Abundance of classical monocytes and their expression of CD127 at day 7 based on CRS incidence (*p < 0.05). **J.** Abundance of transitional monocytes at day 28 based on CRS incidence (*p < 0.05).

At day 7 post-infusion, patients who achieved durable complete remission (CR) at 6 months post-infusion had higher counts of Classical Monocytes than those who later relapsed (p<0.05) (Figure 4D). By day 28, CR patients also exhibited higher neutrophil counts (p<0.05) (Figure 4E).

The composition of CAR+ T-cells infusion products was linked to subsequent CRS development. Patients who developed CRS (grade ≥1, Penn scale(15)) had a higher frequency of CAR+ CD4- MAIT/NKT and CAR+ CD8 Terminal Effector cells in infusion products compared to those without CRS (p<0.05) (Figure 4F). While non-CAR CD8 Terminal Effector cells did not differ between groups, non-CAR CD4- MAIT/NKT cells showed a similar trend (Supplementary Figure 12A).

At day 0 pre-infusion, patients who later developed CRS had fewer plasmablasts, Early and Late NK cells (p<0.05), and pDCs (p<0.01) (Figure 4G) compared to those without CRS. All CRS patients received tocilizumab before day 7. By day 7 post-infusion, tocilizumab-treated patients exhibited significantly higher frequencies of CAR+ CD8 Terminal Effector cells (p<0.05) (Figure 4H), although this trend was not observed in non-CAR CD8 Terminal Effector cells (Supplementary Figure 12B). Additionally, PD-1 expression on CAR+ T-cells at day 7 was lower in tocilizumab-treated patients.

At day 7 post-infusion, patients who experienced CRS had fewer Classical Monocytes and lower expression of IL-7RA (CD127) (p<0.05) on these cells compared to those without CRS (Figure 4I). By day 28, CRS patients had lower counts of Transitional Monocytes (p<0.05) (Figure 4J).

## Discussion

In this study, we investigated the phenotypic composition and dynamics of CD19-targeted CAR-T cells in pediatric patients with R/R BCP-ALL treated with tisagenlecleucel (Novartis, Kymriah)(2, 4, 6). Using mass cytometry, we characterized CAR+ T-cell subsets in infusion products (IP) and tracked their evolution post-infusion, providing insights into treatment response and immune system changes (Figure 5).

**Figure 5.**
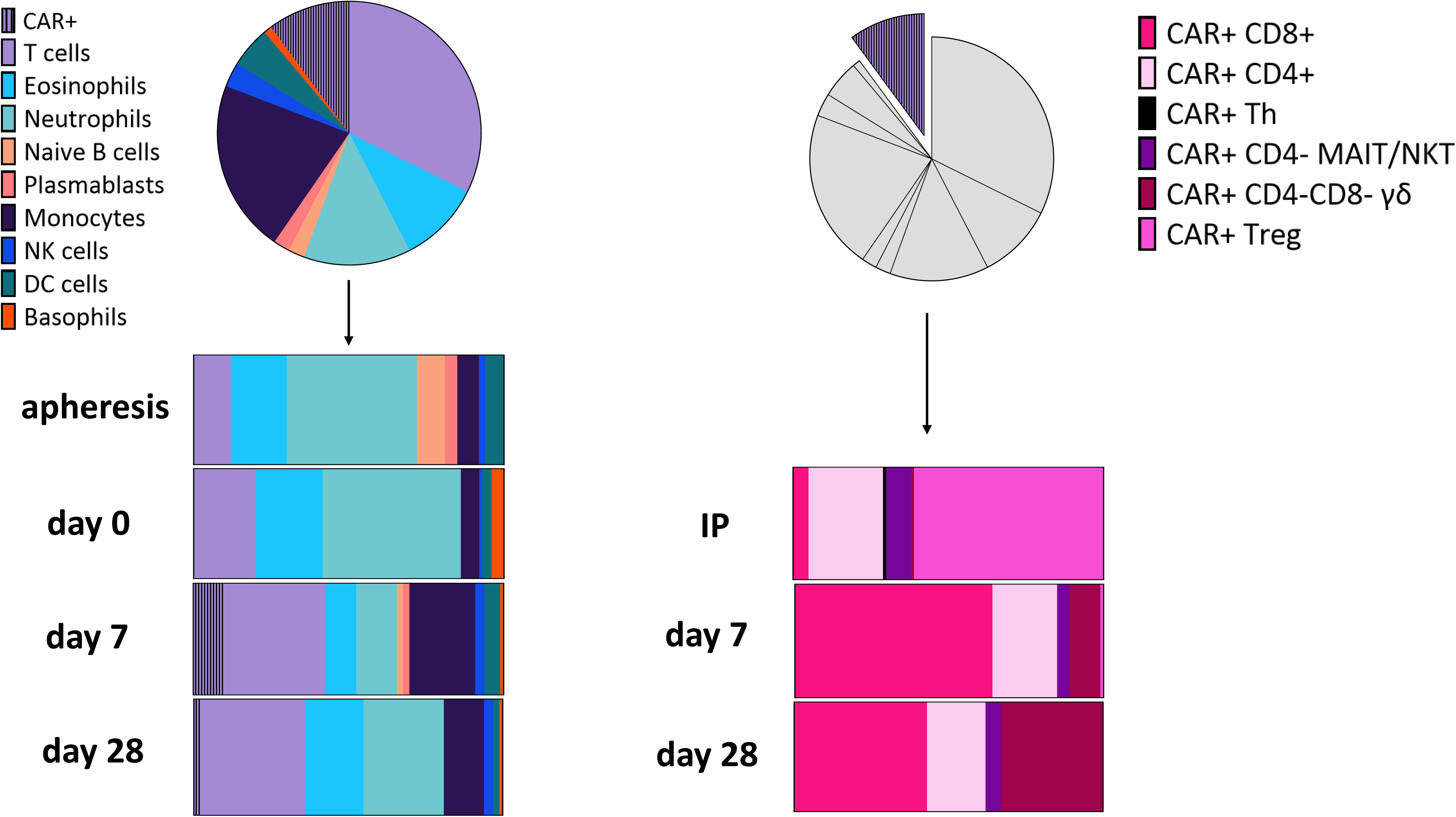
Visual summary of key findings. A graphical overview of CAR+ T-cell subtype dynamics and immune responses throughout CD19 CAR-T therapy. The infusion product (IP) primarily consisted of CAR+ Treg memory cells, with smaller proportions of CD4+ central memory and MAIT/NKT subsets. Following infusion, CAR+ Treg memory cells decline, while CD8+ subsets expand over time. Additionally, monocyte and NK cell levels increase post-infusion, highlighting broader immune activation.

Infusion products were predominantly composed of CAR+ CD4+ T-cells, with a significant fraction exhibiting a central memory phenotype. Notably, CAR+ regulatory T (Treg) memory cells constituted the largest subset, while naïve CD4+ and CD8+ subsets were largely absent. These findings are consistent with previous reports in DLBCL patients treated with tisagenlecleucel, although those studies did not characterize CAR+ Treg memory cells, CD4-CD8- γδ T cells, CD4- MAIT/NKT cells, or T helper cells and were performed among adult patients (9). In contrast to DLBCL, where higher proportions of CD4+ and CD8+ terminal effector cells were observed, these subsets were rare (<2%) in our pediatric BCP-ALL cohort. Despite some interpatient variability, our results align with prior studies in DLBCL(8), suggesting a largely consistent pattern in CAR+ T-cell composition across patients. This consistency is further supported by our analysis of paired infusion products from the same apheresis in three patients, which revealed similar CAR+ T-cell IP profiles within each patient. These findings suggest that while individualized differences exist, the overall phenotypic landscape of infusion products remains relatively stable across patients.

Post-infusion, the proportion of CAR+ Treg memory cells declined by day 7 post-infusion, while CD8+ naïve and effector subsets expanded, indicating a shift toward a more cytotoxic phenotype. This was accompanied by dynamic changes in activation and exhaustion markers, including transient decreases in CD28 and HLA-DR expression, as well as reductions in IL-2Rα (CD25) and CTLA-4 expression at day 7, followed by an increase by day 28. In DLBCL, high CAR+ Treg numbers in infusion products have been associated with tumor growth due to suppression of cytotoxic CAR+ T-cells, correlating with disease progression and reduced neurotoxicity(12). However, in our pediatric BCP-ALL cohort, CAR+ Treg memory cells were absent in PBMCs at days 7 and 28 post-infusion, suggesting a transient effect.

Our data also revealed a post-infusion decline in CAR+ CD4+ Central Memory cells and an expansion of CAR+ CD8+ subsets, consistent with previous studies(23, 24). However, this shift was observed only within 28 days, whereas CAR+ CD4+ have been reported to persist for up to 9-10 years in leukemia patients, correlating with long-term remission. An increase in IL-7Ra (CD127) expression, an activation marker, was observed in our data, particularly in CAR+ CD4+ Central Memory and Effector Memory subsets on day 7 and 28 post-infusion, aligning with previous report(13). Additionally, we did not observe significant changes in PD-1 expression over time. While studies in Hodgkin lymphoma suggest that PD-1+ CD8+ CAR+ T-cells at day 14 correlate with complete response at six months(14), this effect may be disease and age specific.

Beyond CAR+ T-cell dynamics, we observed significant alterations in the broader immune system. As expected, CD19-targeted therapy induced sustained B-cell aplasia(25), reflected in the decline of naïve B cells post-infusion.

Following CAR-T infusion, NK cells expanded and became activated, with increased early and late NK cell subsets by days 7 and 28. The rise in activation markers (CD16, CD161) aligns with reports of enhanced NK cell activity post-infusion(26). Additionally, monocyte dynamics were notable: monocyte numbers increased by day 7 but displayed reduced activity, followed by a decline and subsequent recovery of activity by day 28. Previous studies have linked high monocyte counts in leukapheresis to poor CAR-T cell outcomes(27), low HLA-DR expression to reduced survival(28), and monocyte depletion to improved CAR-T cell manufacturing(29). Our findings extend these observations by revealing dynamic post-infusion monocyte behavior, suggesting a more complex role in therapy response.

While CAR+ T-cell subset composition in infusion products varied among patients, no clear associations were identified between CAR+ T-cell phenotypes and prior treatments (blinatumomab, inotuzumab ozogamicin, hematopoietic stem cell transplantation [HSCT]) or clinical outcomes. Although our assessment of the impact of prior treatment is novel, infusion product phenotypes did not significantly influence outcomes, whereas post-infusion CAR+ T-cell phenotypes in PBMCs might. No consistent associations were observed in tisagenlecleucel studies(9, 10) or in our cohort. However, they have been reports of early memory T cell frequency and Th2 function that correlated with relapse post-infusion(11). In contrast, a higher CD4+:CD8+ ratio in infusion products (varni-cel) has been linked to improved outcomes, potentially due to structural differences in the CAR construct(30).

Higher disease burden is associated with poorer prognosis, including lower event-free survival and increased relapse rates following CD19 CAR-T therapy(31, 32). Consistent with this, patients with a high disease burden in our cohort exhibited altered immune cell composition, including reduced classical monocytes and NK cell subsets in PBMCs at the time of apheresis, which may contribute to adverse outcomes. However, the majority of our patients had a low disease burden at the time of tisagenlecleucel infusion. Eight patients (42%) had measurable disease, as detected by PCR-based minimal residual disease (PCR-MRD) assessment in the bone marrow (≥1×10⁻⁴: positive), but only one patient had a high disease burden (88% bone marrow blasts) prior to infusion. The remaining PCR-MRD-positive patients had less than 1% blasts by morphology, indicating a predominantly low-burden disease cohort.

Our findings challenge the assumption that the absence of absolute neutrophil count (ANC) uniformly enhances CAR-T cell expansion, as previously suggested(33). Instead, we observed that immune recovery, particularly the presence of monocytes and neutrophils, correlates with improved clinical outcomes. This aligns with previous reports linking neutropenia to B-cell recovery(34), suggesting that hematologic reconstitution may play a critical role in sustained remission. These insights underscore the need for further investigation into immune recovery dynamics and their impact on CAR-T therapy efficacy.

The higher frequency of CAR+ CD4- MAIT/NKT and CAR+ CD8+ terminal effector cells in infusion products correlated with CRS development. This aligns with evidence that highly differentiated effector T cells exhibit impaired in vivo function despite strong in vitro activity and that cytokine signaling modifications can influence T cell efficacy(35, 36).

While tocilizumab has proven effective in managing CRS(37), its impact on the immunological profile of CAR+ T-cells remains insufficiently studied. Our findings suggest that baseline immune profiles and monocyte dynamics may serve as predictors of CRS risk in CAR-T therapy. Patients who developed CRS exhibited lower pre-infusion levels of plasmablasts, NK cell, and plasmacytoid dendritic cells (pDCs), indicative of impaired immune regulation(38). By day 7, these patients displayed reduced classical monocyte numbers and IL-7RA expression, suggesting altered immune recovery, potentially influenced by tocilizumab administration(39, 40). By day 28, lower transitional monocyte counts indicated prolonged immune dysregulation(41). These findings suggest that immune profiling could aid in identifying high-risk patients and refining CRS management strategies.

This study has several limitations, including a relatively small cohort, interpatient variability, and retrospective data collection, which may introduce biases. It is also possible that fully elucidating the impacts of patient history on CAR T cell product efficacy, and the impacts of CAR T cell product on patient outcome, requires approaches that can assess the developmental trajectories through lineage tracing(42). Despite these constraints, our study is the first to characterize the phenotypic composition of tisagenlecleucel CAR+ T-cell subsets in pediatric BCP-ALL, revealing a significant number of Tregs in tisa-cell products that do not persist post-infusion in vivo. These findings suggest that the same CD19 CAR-T product may exhibit different behaviors in BCP-ALL compared to lymphoma, as well as in pediatric versus adult patients. Furthermore, our data indicate that the immune system, particularly monocytes and NK cells, play a role in influencing CAR-T treatment outcomes.

Future research should investigate how the immune system shapes CAR-T cell behavior across different malignancies, with a focus on optimizing therapy by targeting key factors such as monocytes and NK cells. A deeper understanding of these immune interactions could help refine CAR-T cell product design, enhance treatment efficacy, and improve outcomes in pediatric BCP-ALL.

## Conclusion

This study provides a detailed characterization of CD19 targeted CAR+ T-cell composition and its evolution in pediatric patients with R/R BCP-ALL treated with tisagenlecleucel. Our findings highlight the predominance of CAR+ CD4+ T cells with a central memory phenotype in infusion products, alongside a significant presence of Treg memory cells. Post-infusion, we observed dynamic shifts in CAR+ T-cell subsets, including a decline in Treg memory cells and an expansion of CD8+ effector populations. These changes were associated with immune system alterations and clinical responses, particularly CRS. By mapping CAR+ T-cell dynamics from infusion to post-treatment, this study provides valuable insights into their role in therapy response. These findings may help to refining CAR-T product design and improving patient outcomes in pediatric BCP-ALL.

## Declarations

### Ethics approval and consent to participate

This study involves human participants and was approved by the Human Research Ethics Committee of the Medical University of Lodz (approval no. RNN/93/22/KE). Participants gave informed consent to participate in the study before taking part, according to the Helsinki declaration and its subsequent amendments. No personally identifiable information was included in the paper.

### Consent for publication

All authors authorized and granted full consent to the corresponding author of the manuscript (AO) to enter into publishing agreement.

### Availability of data and material

All data relevant to the study are included in the article or uploaded as supplementary information.

### Authors’ contributions

AO – conceived the study and secured funding. KLD, WM conceptualized the study and supervised the project. AO, BP, NCP, JCC, SJ, MP, MPV developed methods. AO, BP, NCP and MM performed all experiments. AO and AK developed the software. AO, AK and JCC performed formal data analysis for all of the data generated. PM, MRP, MMS, KK, JS treated patients and/or acquired clinical samples and data. AO, MP, KLD and WM interpreted the results and wrote the first draft of the paper. All authors critically reviewed the manuscript.

## List of Abbreviations

CAR-T: chimeric antigen receptor T-cell
BCP-ALL: B-cell precursor acute lymphoblastic leukemia
DLBCL: diffuse large B-cell lymphoma
FDA: U.S. Food and Drug Administration
EMA: European Medicines Agency
PBMC: peripheral blood mononuclear cell
CRS: cytokine release syndrome
EFS: event-free survival
OS: overall survival
IP: infusion product
tisa-cel: Tisagenlecleucel
PCR-MRD: PCR-based minimal residual disease
allo-HSCT: allogeneic hematopoietic stem cell transplantation
R/R BCP-ALL: relapsed/refractory B-cell precursor acute lymphoblastic leukemia
pDCs: plasmacytoid dendritic cells
mDCs: myeloid dendritic cells
ANC: absolute neutrophil count
Treg: regulatory T

