## Supplemental materials for "Single-cell profiling of CAR-T CD19 cell phenotypes and immune system dynamics in pediatric BCP-ALL"

**Supplementary Table 1.**

Single cell (Mass cytometry) analysis was performed using Maxpar Direct Immune Profiling Assay, that has 30-marker panel showed in table. Additionally three immune checkpoints antigens (PD-1, PD-L1, CTLA-4) and antigen for CAR were added. Table presents antibodies combined to heavy metals.

| **Target** | **Metal** |
|---|---|
| Anti-human CD45 | 89Y |
| Live/dead 103Rh-Intercalator (500 µM) | 103Rh |
| Anti-human CD196/CCR6 | 141Pr |
| Anti-human CD123 | 143Nd |
| Anti-human CD19 | 144Nd |
| Anti-human CD4 | 145Nd |
| Anti-human CD8a | 146Nd |
| Anti-human CD11c | 147Sm |
| Anti-human CD16 | 148Nd |
| Anti-human CD45RO | 149Sm |
| Anti-human CD45RA | 150Nd |
| Anti-human CD161 | 151Eu |
| Anti-human CD194/CCR4 | 152Sm |
| Anti-human CD25 | 153Eu |
| Anti-human CD27 | 154Sm |
| Anti-human CD57 | 155Gd |
| Anti-human CD183/CXCR3 | 156Gd |
| Anti-human CD185/CXCR5 | 158Gd |
| **Monoclonal Anti-FMC63 scFv Antibody, Mouse IgG1 (Y45)** | **159 Tb** |
| Anti-human CD28 | 160Gd |
| Anti-human CD38 | 161Dy |
| Anti-human CD56/NCAM | 163Dy |
| Anti-human TCRgd | 164Dy |

| | |
|---|---|
| **Anti-Human CD279/PD-1 (EH12.2H7)** | **165Ho** |
| Anti-human CD294 | 166Er |
| Anti-human CD197/CCR7 | 167Er |
| Anti-human CD14 | 168Er |
| **Anti-human CTLA-4** | **169Tm** |
| Anti-human CD3 | 170Er |
| Anti-human CD20 | 171Yb |
| Anti-human CD66b | 172Yb |
| Anti-human HLA-DR | 173Yb |
| Anti-human IgD | 174Yb |
| **Anti-Human CD274/PDL1 (29E.2A3)** | **175Lu** |
| Anti-human CD127 | 176Yb |

**Supplementary Figure 1.**

**(a)**

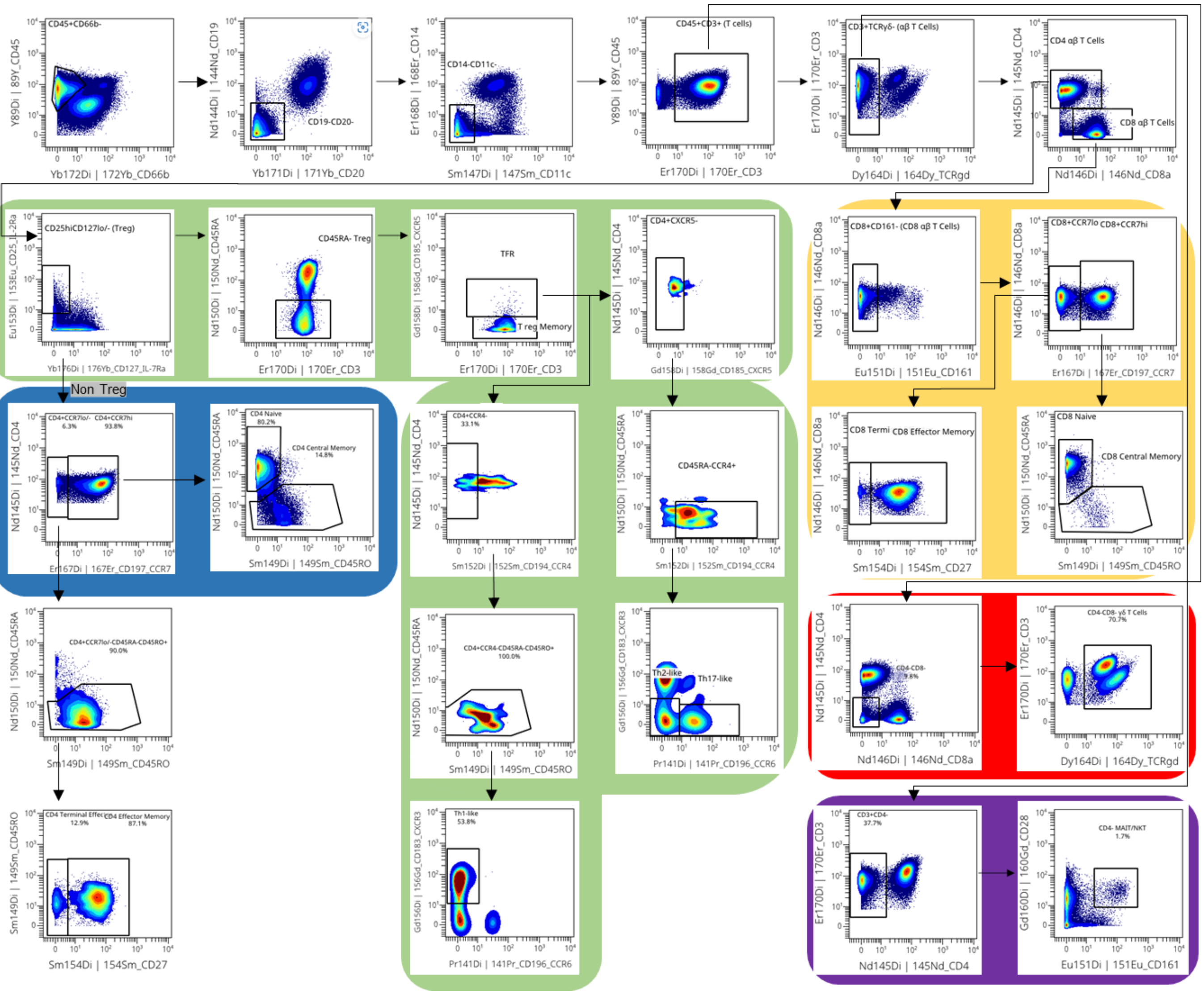

(b)

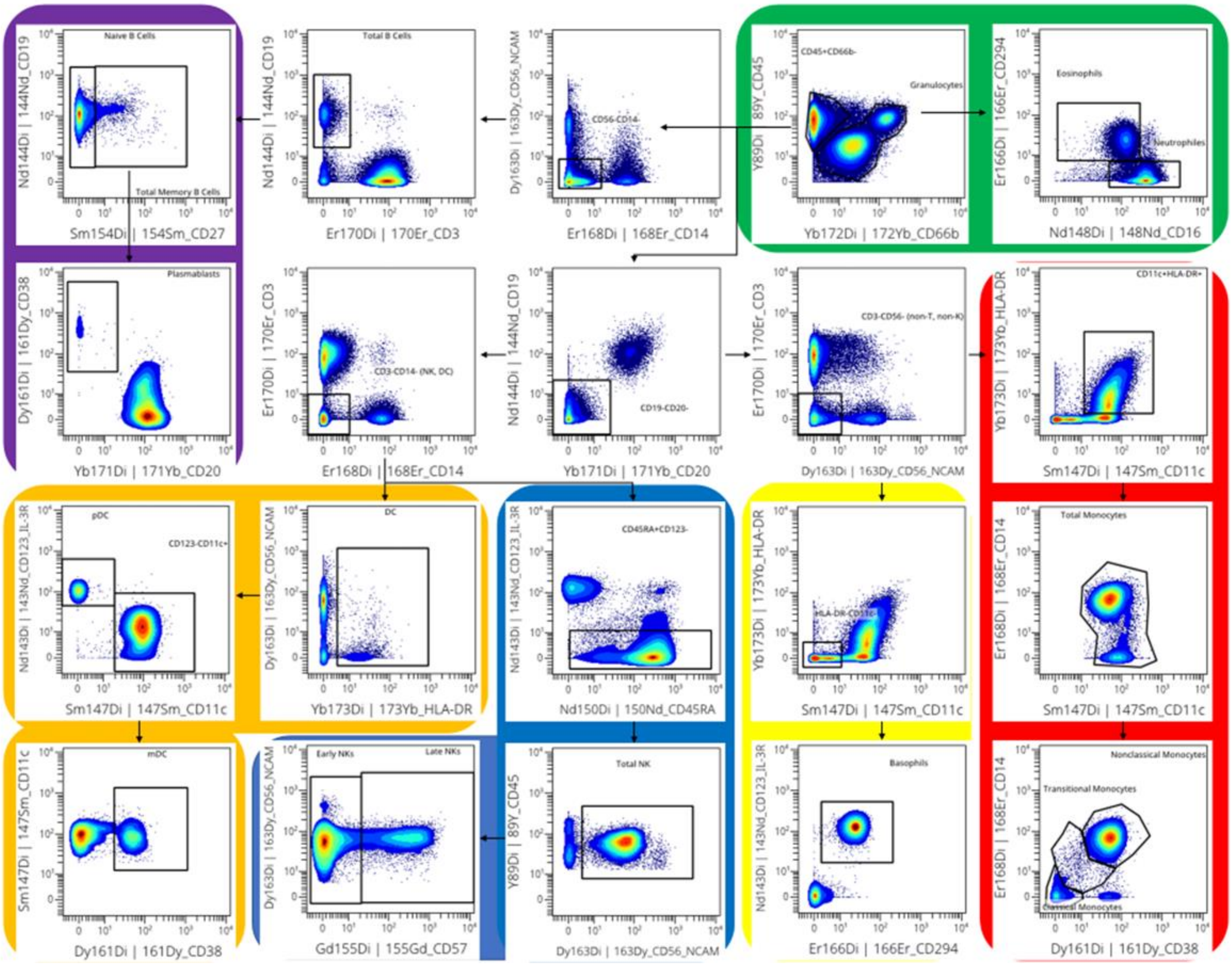


**Supplementary Figure 1. Gating strategy.**

**(a) T cells gating strategy.**

Flow plot showing gating strategy for T cells from Human healthy peripheral blood (4 PBMC from healthy children, ~300,000 cells per sample). Gating and sub sequential analysis was done using OMIQ.

Color green highlights gating for T-reg memory and Th-like 1,2,17; Blue – CD4+; Yellow – CD8+; Red – CD4-CD8- γδ T cell, Purple – CD4- MAIT/NKT.

**(b) Other cells gating strategy.**

Flow plot showing gating strategy for cells from Human healthy peripheral blood (4 PBMC from healthy children, ~300,000 cells per sample). Gating and sub sequential analysis was done using OMIQ.

Color green highlights gating for neutrophils and eosinophils; Red – monocytes; Yellow – basophils; Blue – NK cells; Orange – DC; Purple – B cells.

**Supplementary Figure 2.**

**(a)**

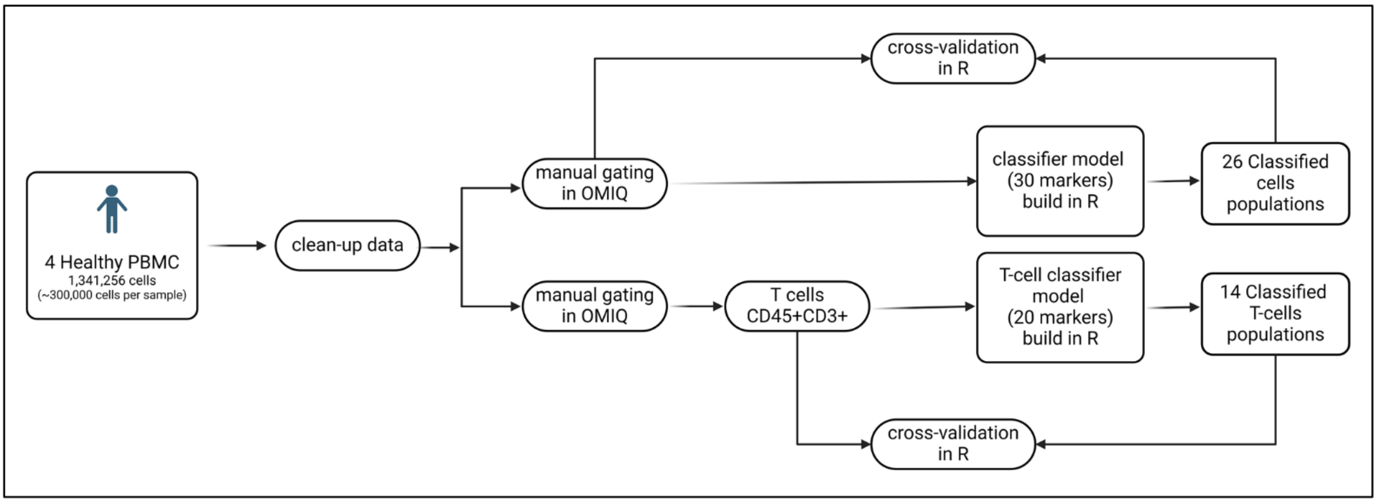


**(b)**

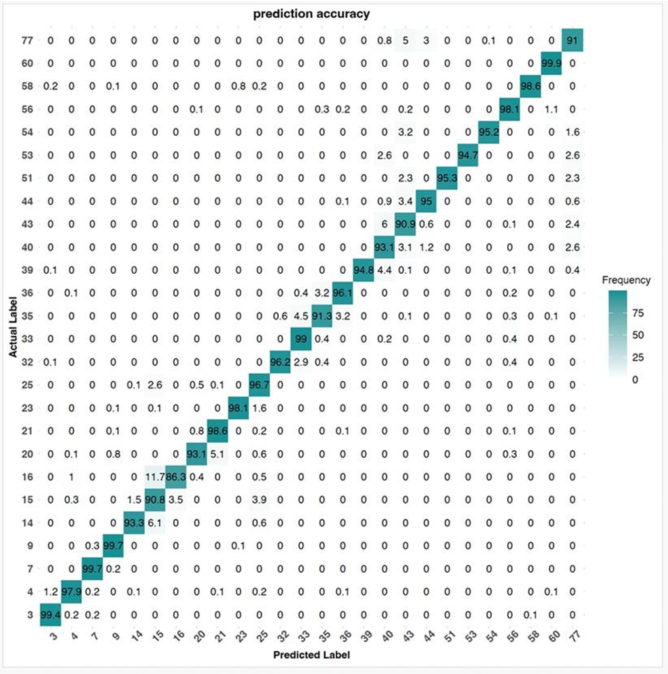

(c)

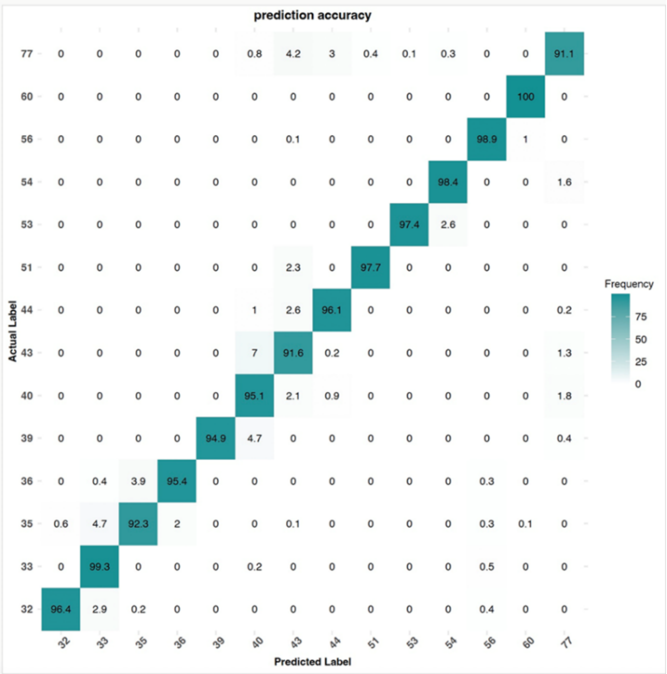


(d)

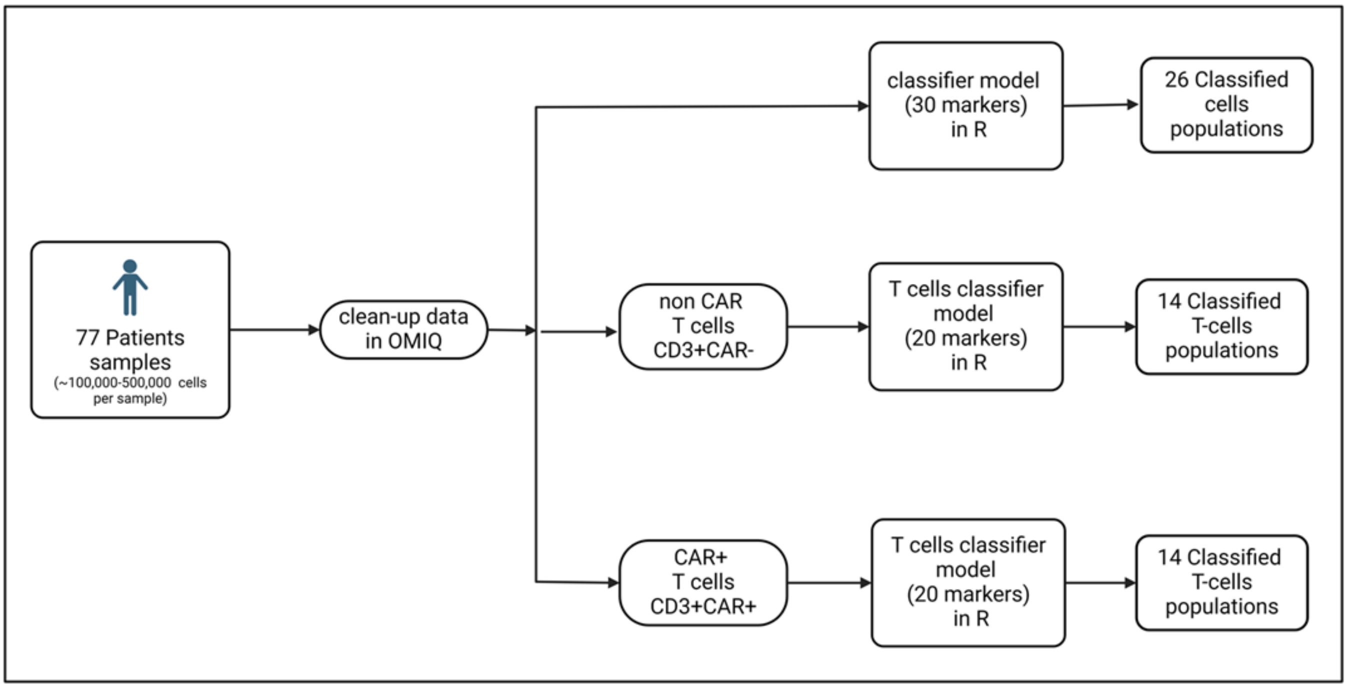


**Supplementary Figure 2. Classification Model for Cell Populations and T-Cell Subpopulations.**

**(a) Methodology Overview.**

Schematic representation of the classification model constructed in R for identifying all cell populations and specifically for T-cell populations. The diagram outlines the workflow used to classify cell populations, followed by a cross-validation process to evaluate model performance. Diagram created with BioRender.com**.**

**(b,c) Heatmap demonstrating accuracy of classification in healthy controls, as a cross validation for classifier method for (b) all populations and (c) T cells populations.**

77- Treg memory; 60 – CD4-CD8- γδ T cells; 58 – Basophils; 56 – CD4- MAIT/NKT; 54 – Th17-like; 53 – Th2-like; 51 – Th1-like; 44 – CD4 Terminal Effector; 43 – CD4 Effector Memory; 40 – CD4 Central Memory; 39 – CD4 Naïve; 36 – CD8 Terminal Effector; 35 – CD8 Effector Memory; 33 – CD8 Central

Memory; 32 – CD8 Naïve; 25 – mDC; 23 – pDC; 21 – Late NKs; 20 – Eayly NKs; 16 – Nonclassical Monocytes; 15 – Transitional Monocytes; 14 – Classical Monocytes; 9 – Plasmablasts; 7 – Naive B cells; 4 – Neutrophils; 3 – Eosinophils.

**(d) Application of Supervised Classification**

The diagram illustrates the application of a supervised classification method. The classifier model built in R is used sequentially: first to classify all cell populations, then to classify non-CAR T cells, and finally to classify CAR+ T-cells. Diagram created with BioRender.com.

**Supplementary Figure 3.**

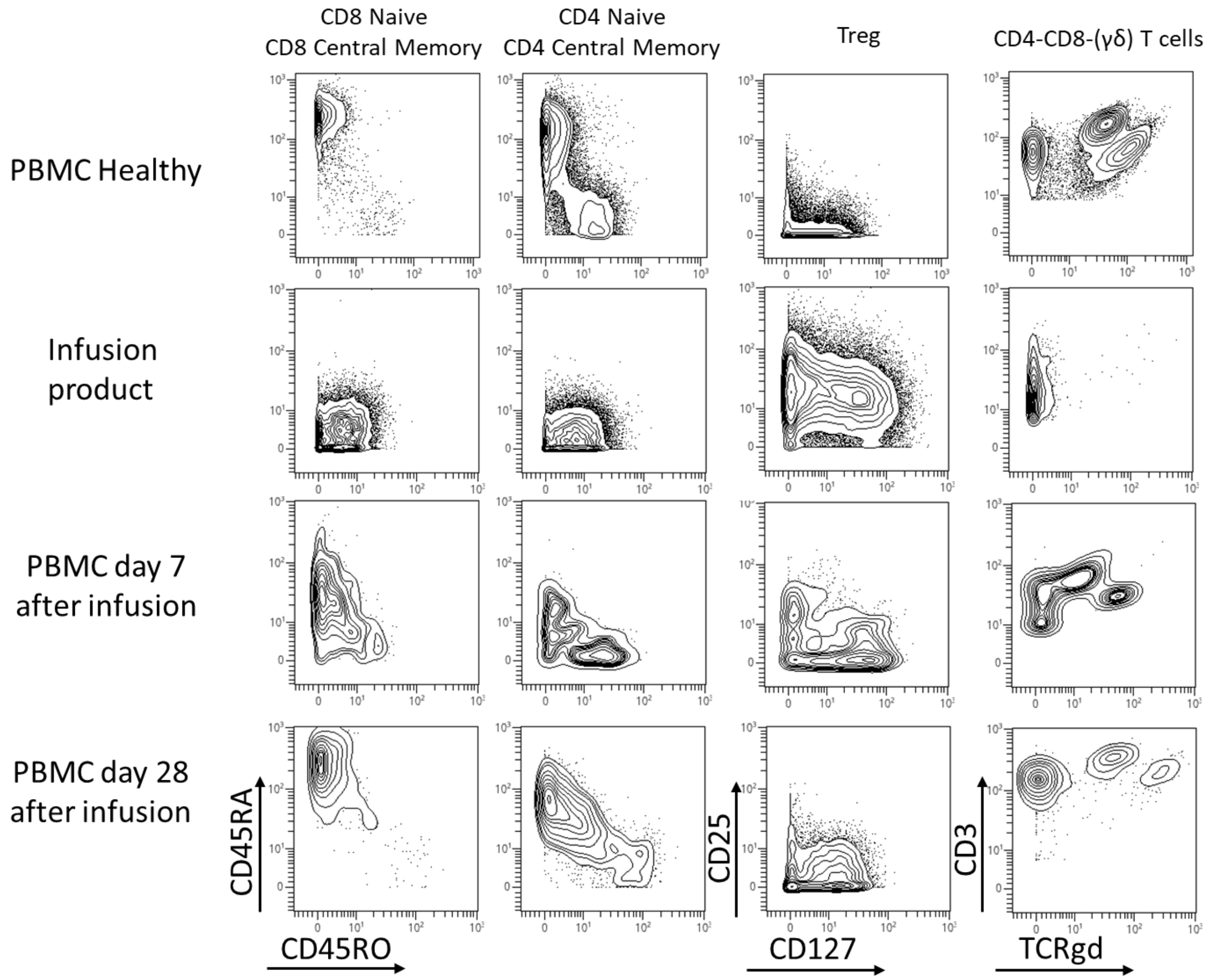


**Supplementary Figure 3.**
**Mass cytometry analysis of CD8 Naïve/CD8 Central Memory, CD4 Naïve/CD4 Central Memory, Treg and CD4-CD8- γδ T cells subpopulations of T cells.**
Samples from three representative patients with BCP-ALL and one healthy donor.

**Supplementary Figure 4.**

**(a)**

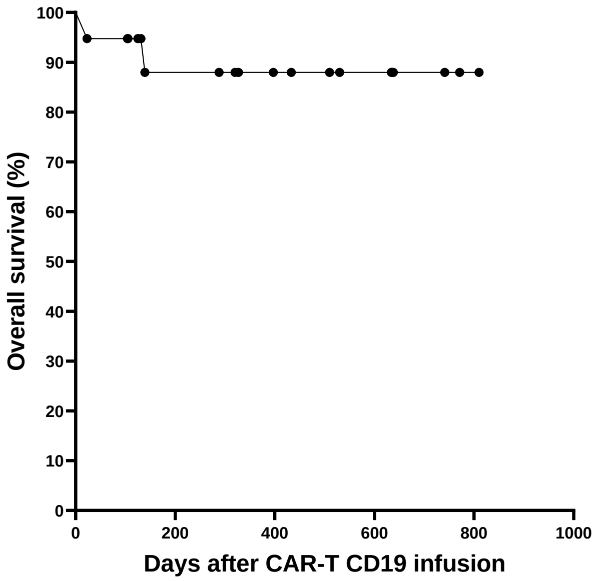


**(b)**

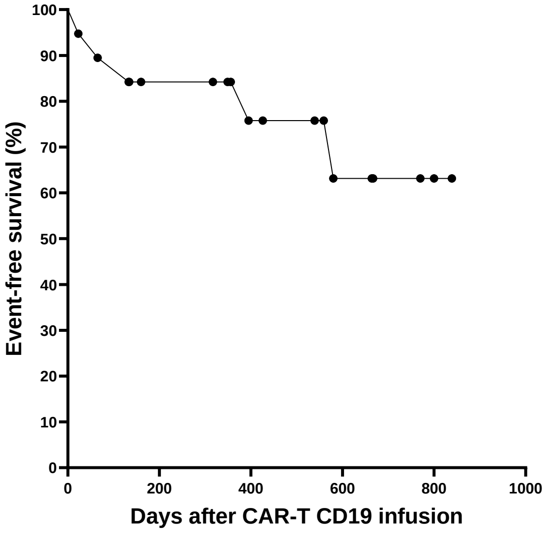


**Supplementary Figure 4. Patients' survival.**

Kaplan–Meier curve showing OS (a) and EFS (b). Patients have been observed from 0.5 year – 2 year (median 1 year). All patients have been observed for at least 6 months. 6-months OS was 87.97%, 6-months EFS was 74.56%.

**Supplementary Figure 5.**

**(a)**

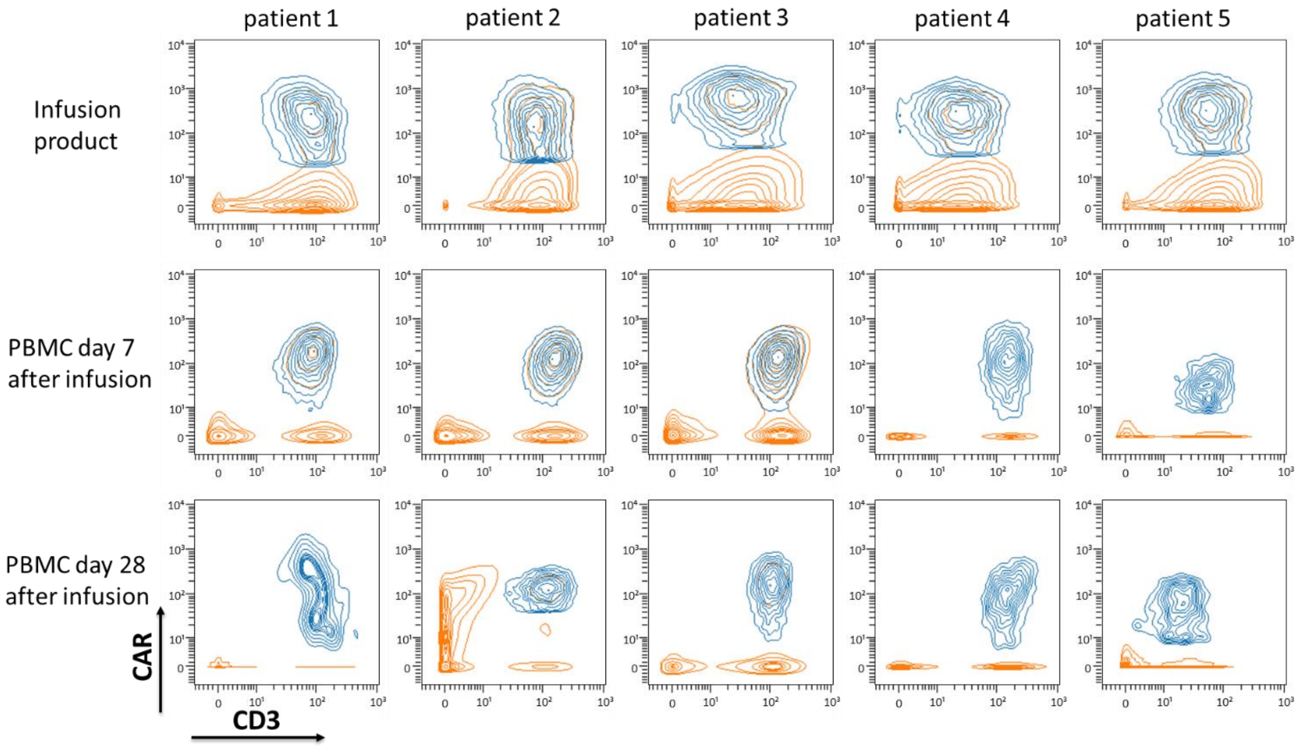


**(b)**

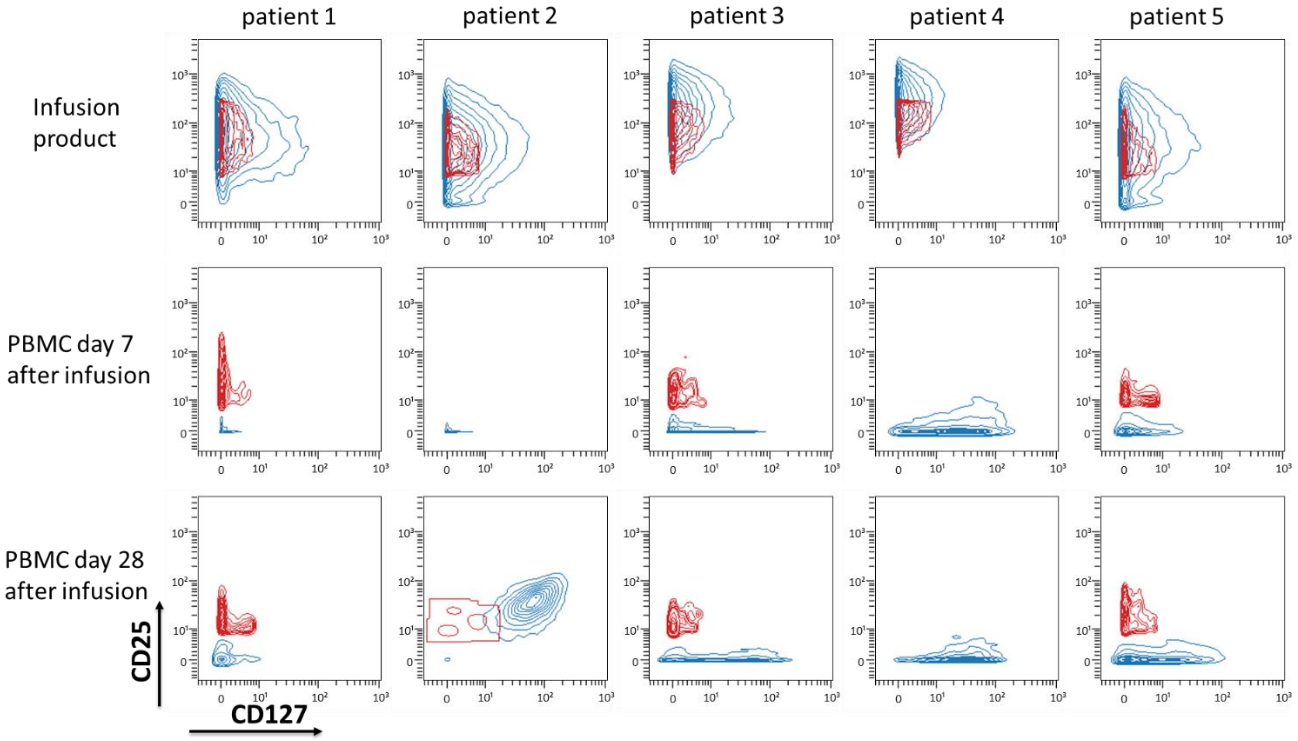

**(c)**

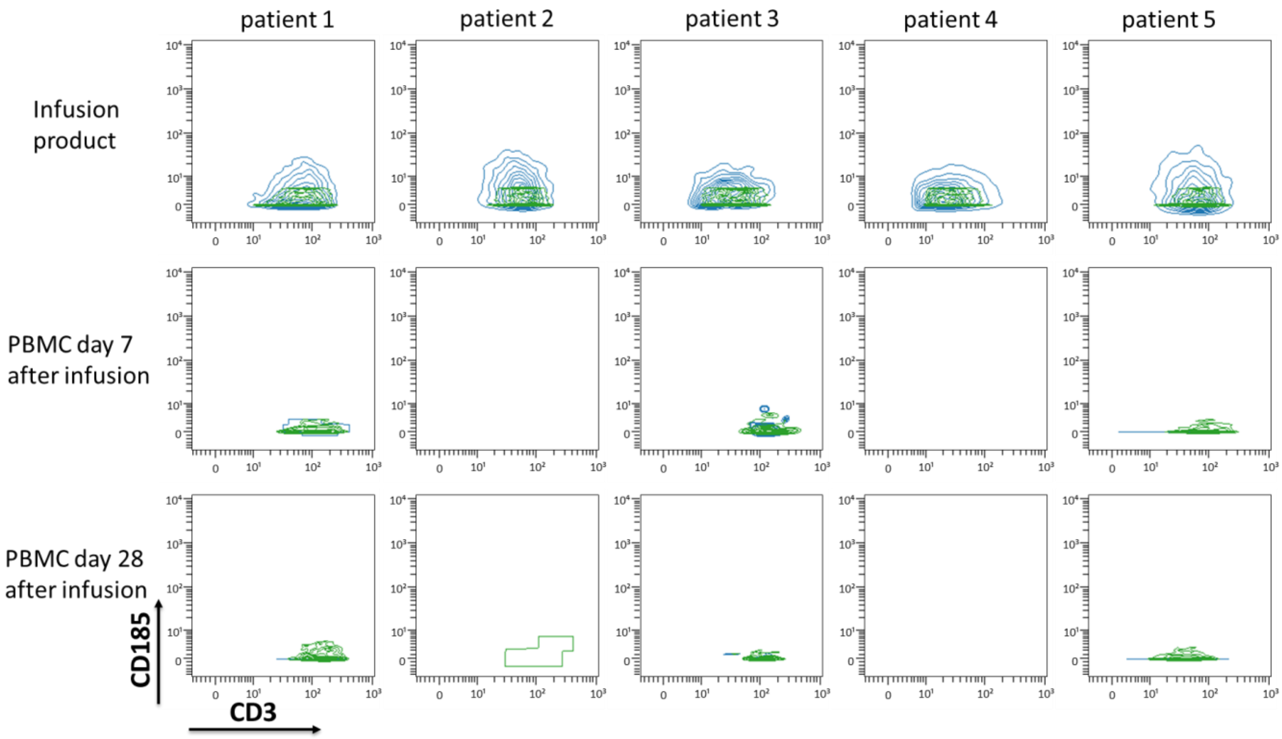


**Supplementary Figure 5. Manual gating of CAR+ T-cells (a), Treg (b), and Treg memory (c) as validation of the applied method.**

Graphs (a) and (b) were generated using unfiltered cells, whereas graph (c) was filtered for Treg cells. Samples were collected from five representative patients with BCP-ALL in IP, followed by PBMC analysis on days 7 and 28. Color coding: orange – unfiltered cells, blue – CAR+ T-cells, red – Treg cells, green – Treg memory cells.

**Supplementary Figure 6.**

**(a)**

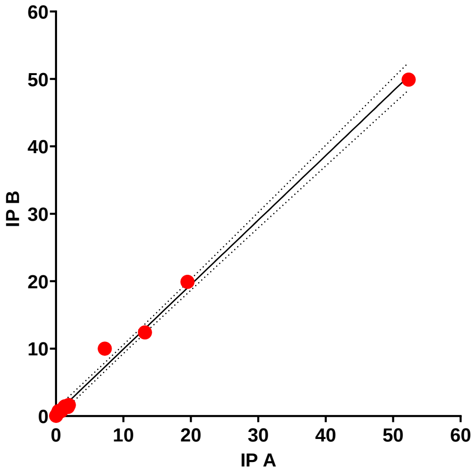


**(b)**

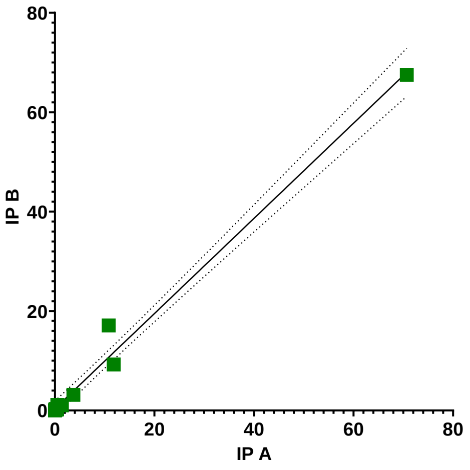


**(c)**

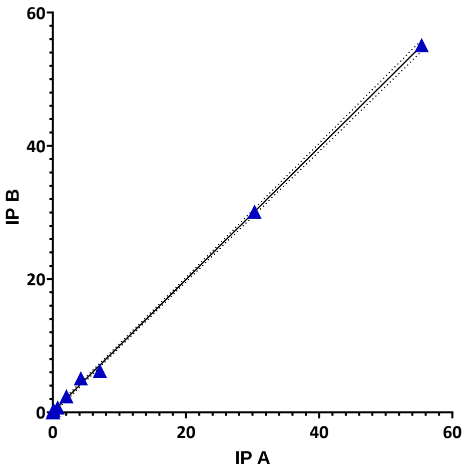


**Supplementary Figure 6. Comparative analysis of two CAR-T CD19 infusion products (IP A and IP B) administered to three patients (Patients 6 (a), 12 (b), and 18 (c)) reveals patient-specific variability in CAR+ T-cell subpopulations.**

Simple linear regression was calculated, with R^2=0.9957(a), R^2=0.9881(b), R^2=0.9995(c) and p value <0.0001.

**Supplementary Figure 7.**

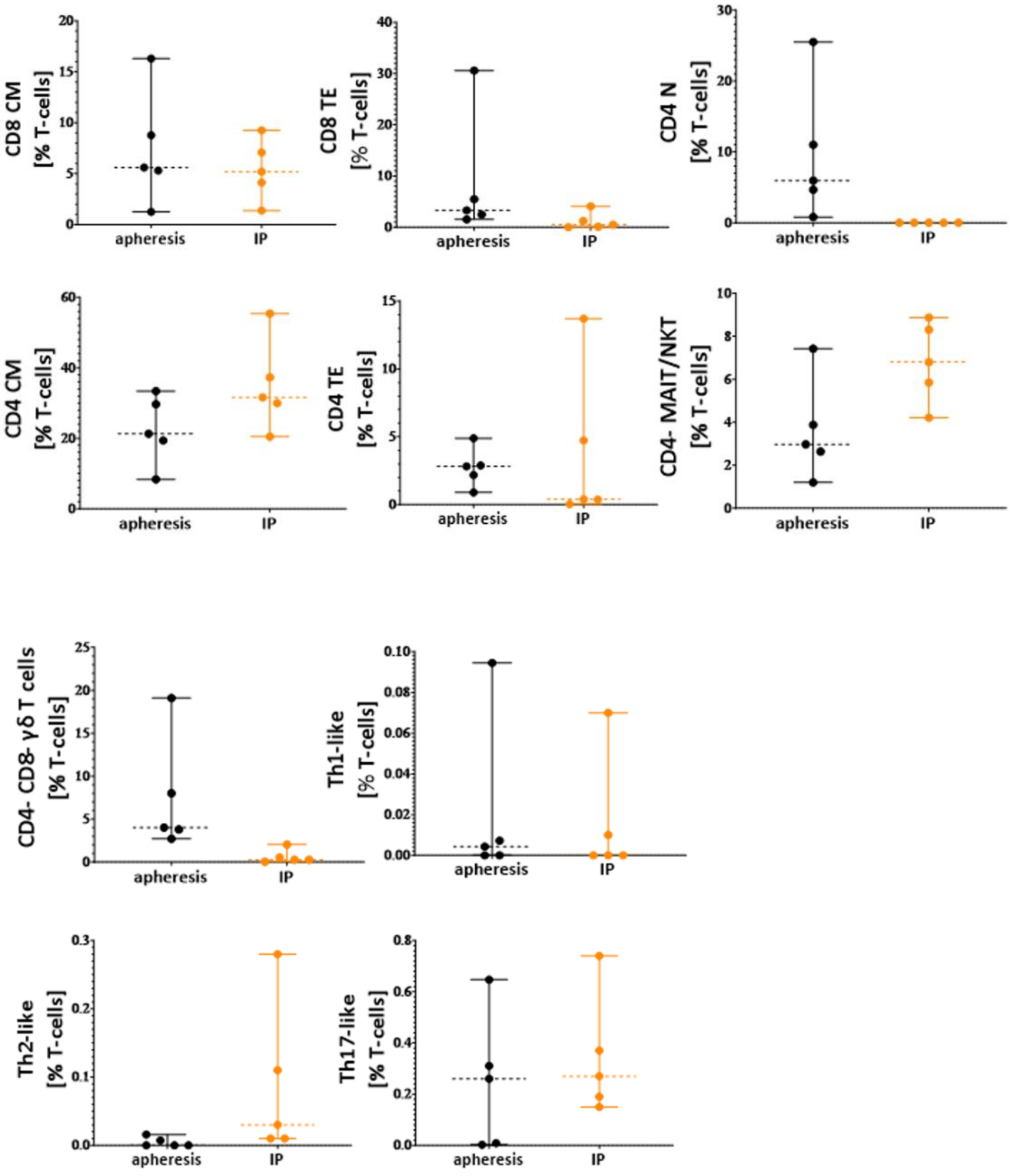


**Supplementary Figure 7. Mass cytometry analysis of CAR+ T-cell subpopulations during CAR-T CD19 therapy.**

Comparison of phenotype of T cell subpopulations at the time of apheresis and CAR+ T-cells from infusion products.

Violin plots showing the comparison of CD8 Central Memory, CD8 Terminal Effector, CD4 Naïve, CD4 Central Memory, CD4 Terminal Effector, CD4- MAIT/NKT, CD4-CD8- γδ T cells, Th1-like, Th2-like and Th17-like between PBMC at the time of apheresis and matching infusion products for 5.

P values were calculated applying the paired t-test.

**Supplementary Figure 8.**

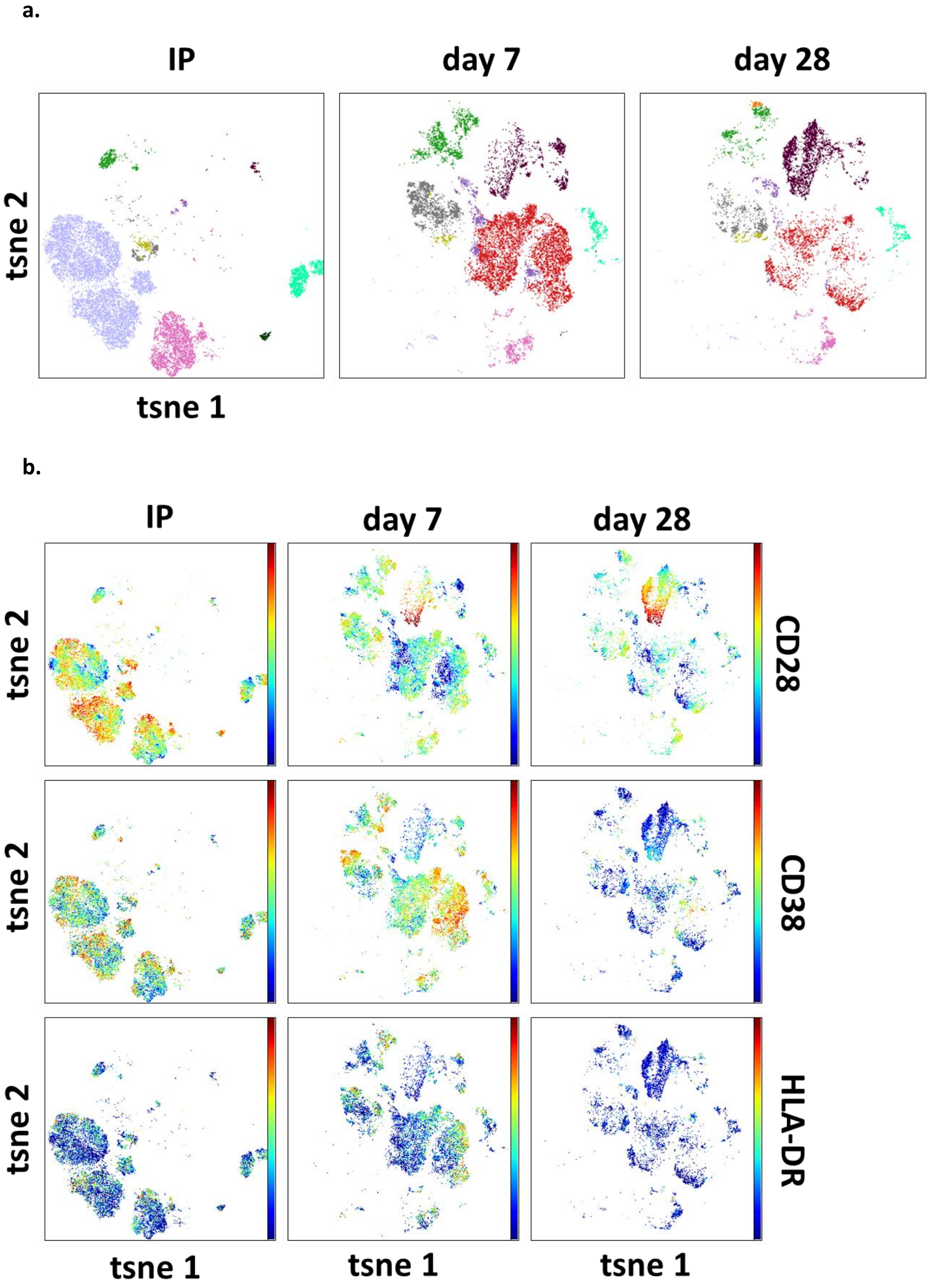

**c.**

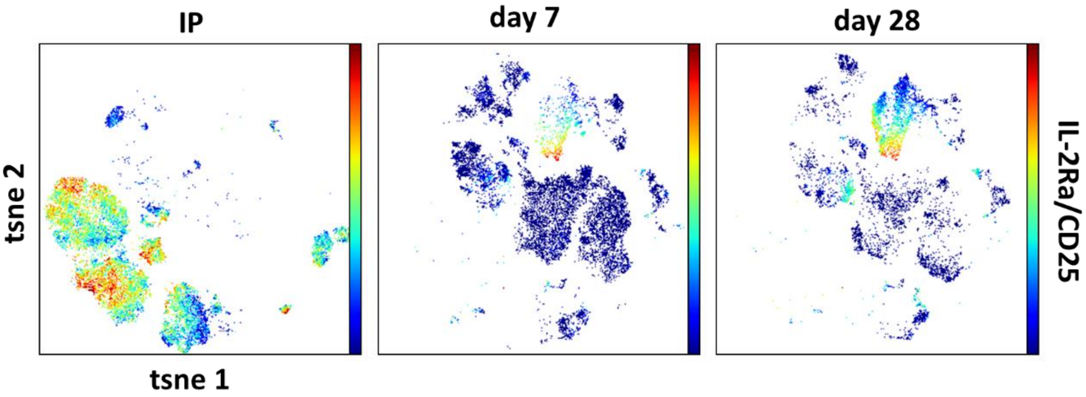


**d.**

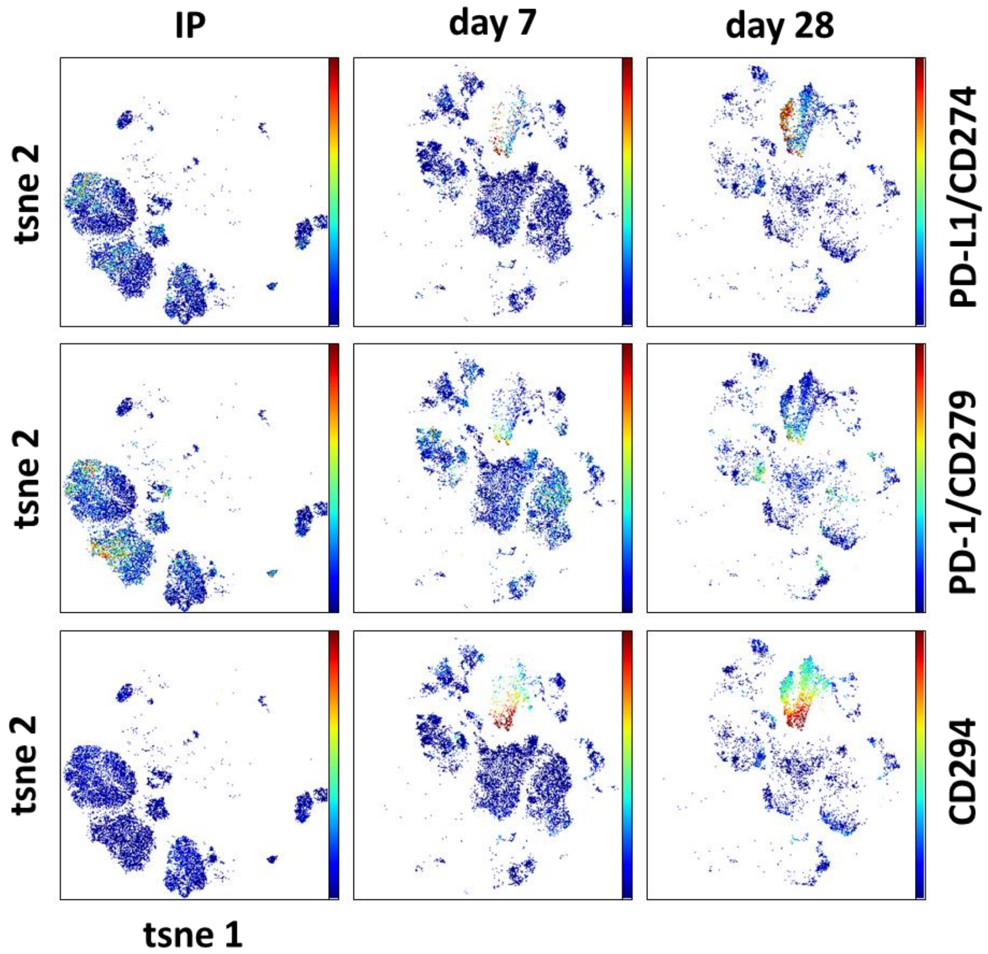


**Supplementary Figure 8.**

**Expression of exhaustion and activation markers on CAR+ T-cells during CD19 CAR-T therapy.**

tSNE analysis of CAR+ T-cells at different time points (IP, day 7, and day 28) reveals distinct CAR+ T-cell subsets (a). tSNE plots further illustrate the expression of activation markers (CD28, CD38, HLA-DR) within these subsets (b), the expression of IL-2RA (c), and exhaustion markers (PD-L1, PD-1, CD294) (d).

**Supplementary Figure 9**

(a)

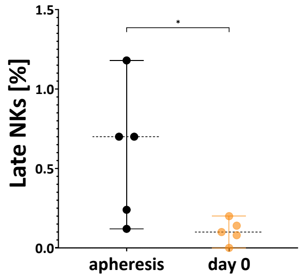


(b)

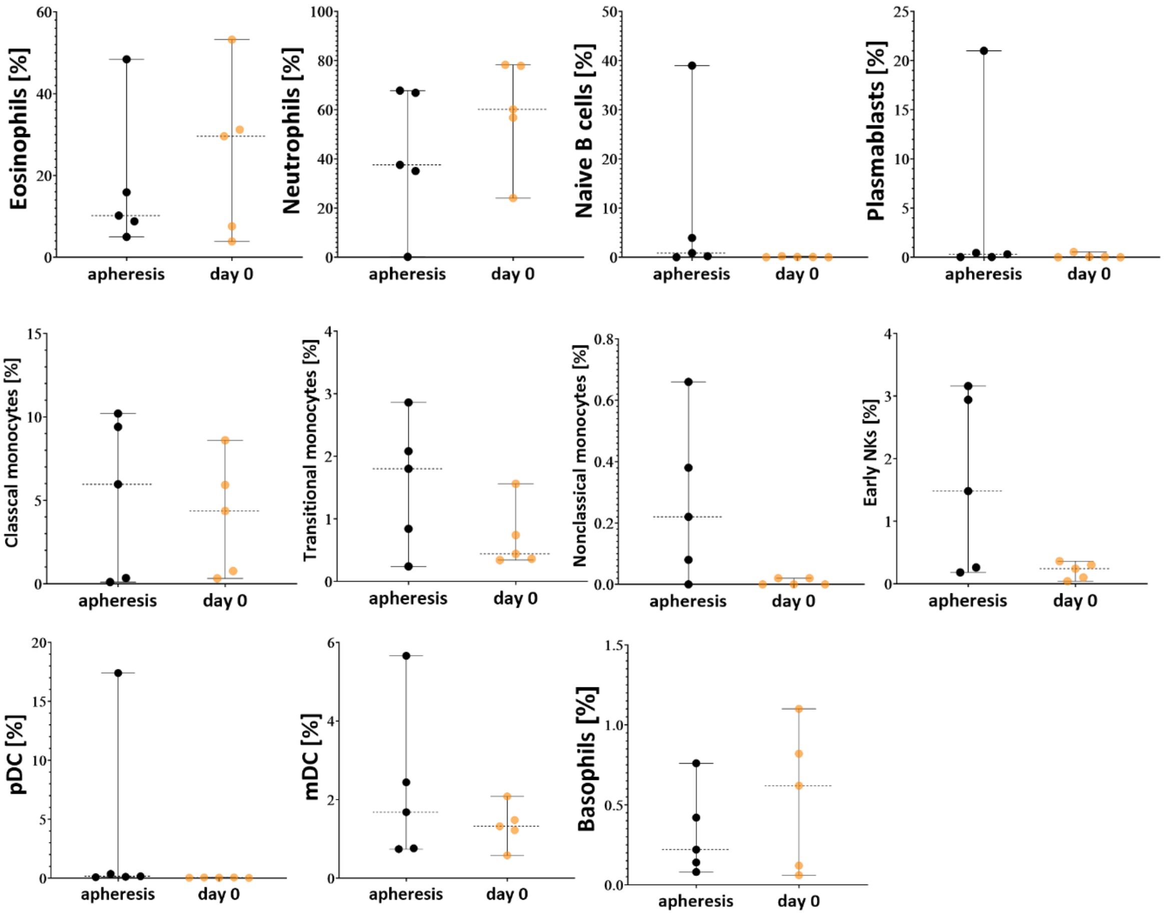

(c)

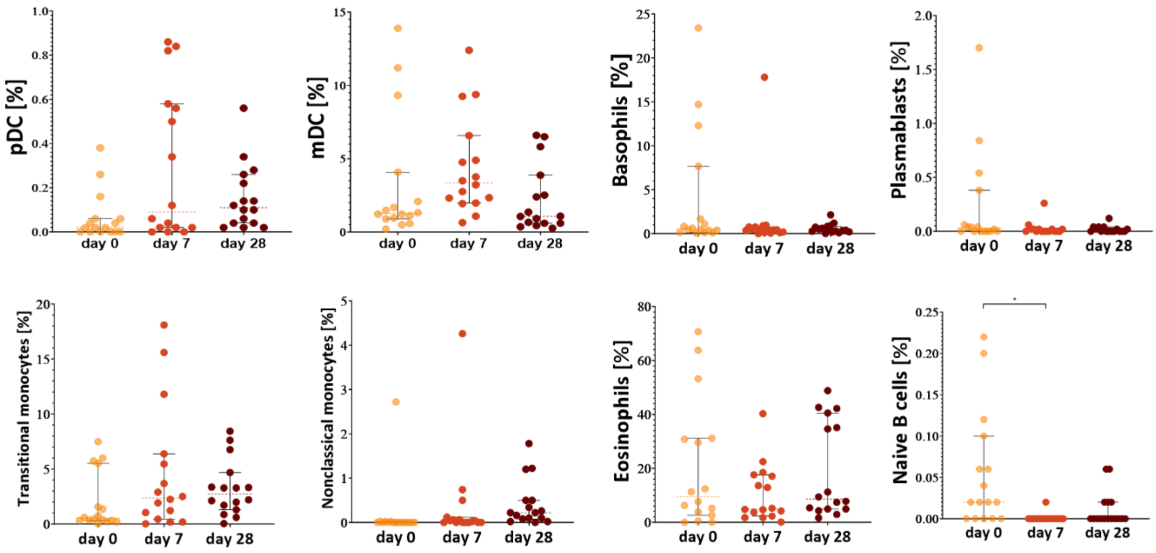


**Supplementary Figure 9. Mass cytometry analysis of transition of cell subpopulations during CAR-T CD19 therapy.**

- **(a) Late NK cells in apheresis samples compared to day 0 (*p < 0.05).**
- **(b) Comparison of cell populations in the PBMC of BCP-ALL patients during CAR-T CD19 therapy between apheresis and day 0 (before CAR-T CD19 infusion).**

Violin plots showing the comparison of cells populations for 5 patients (all matching samples). P values were calculated applying the paired t-test.

- **(c) Comparison of cell populations in the PBMC of BCP-ALL patients during CAR-T CD19 therapy between day 0 (before CAR-T CD19 infusion) and day 7 and day 28 after CAR-T CD19 infusion.**

Violin plots showing the comparison of cells populations for 16 patients (all matching samples). P values were calculated applying the paired t-test (*p<0.05).

**Supplementary Figure 10.**

**(a)**

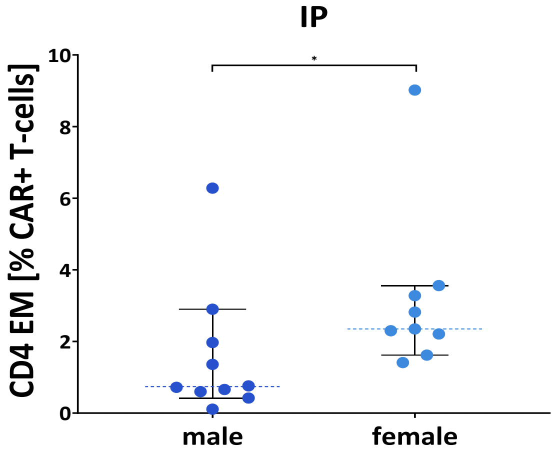


**(b)**

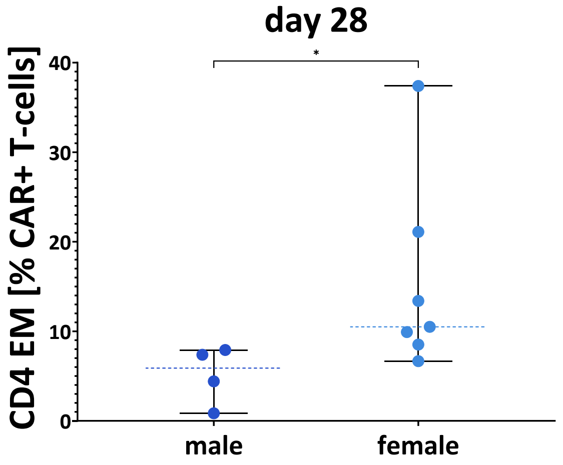


**Supplementary Figure 10.**

**Higher percentage among CAR+ T-cells of CAR+ CD4 Effector Memory in female vs male, in the CAR-T CD19 infusion products, and following in the PBMC at the day 28.**

Violin plots showing difference in the percentage of CAR+ CD4 Effector memory within the CAR-T CD19 infusion products (a) and following 28 days in PBMC (b) between female and male. P values were calculated applying the Mann Whitney test. *p < 0.05; **p < 0.01.

**Supplementary Figure 11.**

(a)

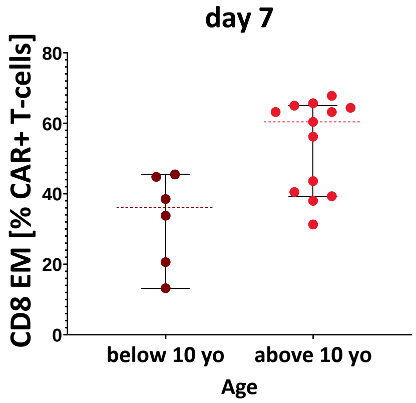


(b)

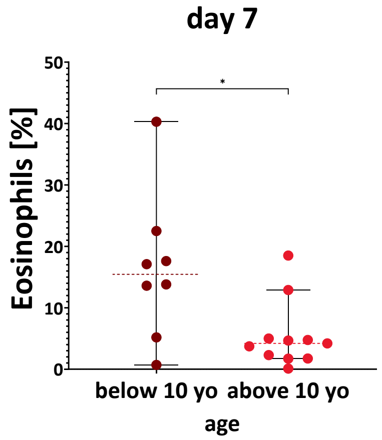


(c)

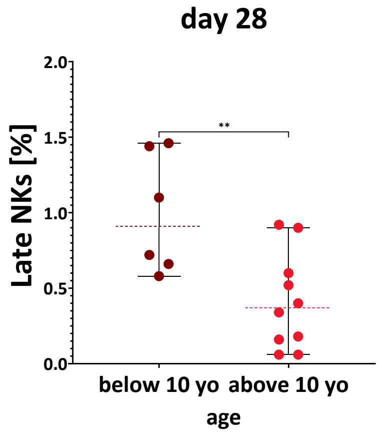

**Supplementary Figure 11. Age-Associated Differences in CAR+ CD8 Effector Memory, Eosinophils, and Late NK Cells in PBMCs Post-Infusion.**

**(a) A higher proportion of CAR+ CD8 Effector Memory cells among CAR+ T-cells is observed in older patients (≥10 years) compared to younger patients (<10 years) in PBMCs on day 7 post-infusion.**

A significant difference in CAR+ CD8 Effector Memory cells was detected between these age groups. Statistical significance was determined using the Mann-Whitney U test (*$p < 0.05$).

**(b) Differential eosinophil counts in PBMCs based on patient age.**

Violin plots illustrate higher eosinophil counts in PBMCs of patients younger than 10 years compared to those aged 10 years or older. A statistically significant difference in eosinophil counts was observed at day 7 post-infusion. Statistical significance was determined using the Mann-Whitney U test (*$p < 0.05$, **$p < 0.01$).

**(c) Differential late NK cell counts in PBMCs based on patient age.**

Violin plots show higher late NK cell counts in PBMCs of patients younger than 10 years compared to those aged 10 years or older. A statistically significant difference in late NK cell counts was observed at day 28 post-infusion. Statistical significance was determined using the Mann-Whitney U test (*$p < 0.05$, **$p < 0.01$).

**Supplementary Figure 12.**

**(a)**

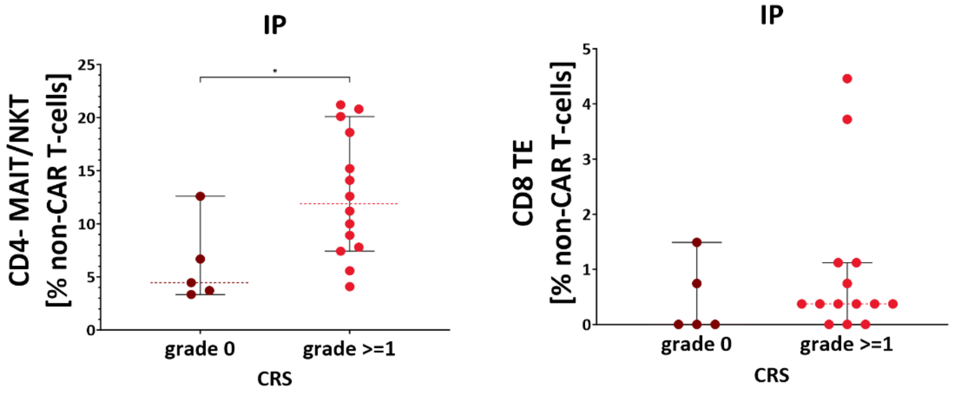


**(b)**

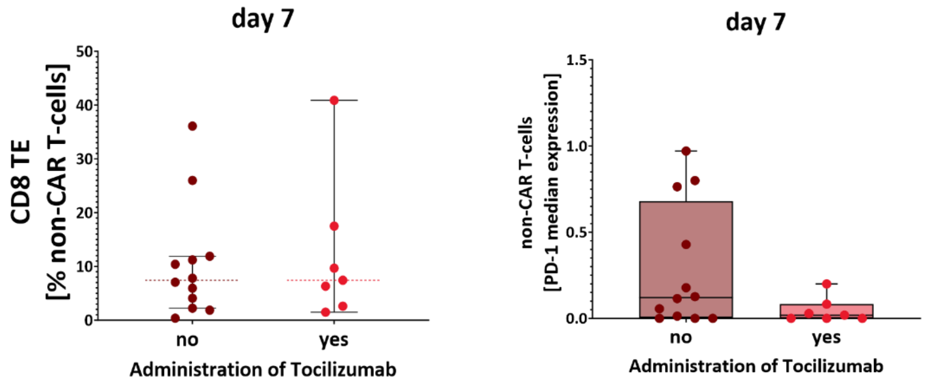


**Supplementary Figure 12.**

**Association of CRS with non-CAR subpopulations during CD19 CAR-T therapy.**

**(a) Differences in non-CAR cells in infusion products.**

Patients who developed CRS (grade ≥1) exhibited significantly higher levels of non-CAR $CD4^-$ MAIT/NKT cells compared to asymptomatic patients (grade 0). However, differences in non-CAR CD8 Terminal Effector cells were not statistically significant ($p = 0.4039$). P-values were calculated using the Mann-Whitney U test (*$p < 0.05$).

**(b) Impact of tocilizumab on non-CAR cells at day 7 post-infusion.**

At day 7 post-infusion, no significant differences in the non-CAR CD8 Terminal Effector population were observed between patients who received tocilizumab for CRS and those who did not. All patients received tocilizumab prior to day 7 post-infusion. Additionally, PD-1 expression on non-CAR cells did not significantly differ between patients who received tocilizumab and those who did not. P-values were calculated using the Mann-Whitney U test.
